# Development of a Coupled Simulation Toolkit for Computational Radiation Biology Based on Geant4 and CompuCell3D

**DOI:** 10.1101/2020.04.09.034926

**Authors:** Ruirui Liu, Kathryn A. Higley, Maciej H. Swat, Mark A. J Chaplain, Gibin G. Powathil, James A. Glazier

## Abstract

Understanding and designing clinical radiation therapy is one of the most important areas of state-of-the-art oncological treatment regimens. Decades of research have gone into developing sophisticated treatment devices and optimization protocols for schedules and dosages. In this paper, we presented a comprehensive computational platform that facilitates building of the sophisticated multi-cell-based model of how radiation affects the biology of living tissue. We designed and implemented a coupled simulation method, including a radiation transport model, and a cell biology model, to simulate the tumor response after irradiation. The radiation transport simulation was implemented through Geant4 which is an open-source Monte Carlo simulation platform that provides many flexibilities for users, as well as low energy DNA damage simulation physics, Geant4-DNA. The cell biology simulation was implemented using CompuCell3D (CC3D) which is a cell biology simulation platform. In order to couple Geant4 solver with CC3D, we developed a “bridging” module that extracts tumor cellular geometry of the CC3D simulation (including specification of the individual cells) and ported it to the Geant4 for radiation transport simulation. The cell dose and cell DNA damage distribution in multicellular system were obtained using Geant4. The tumor response was simulated using cell-based tissue models based on CC3D. By merging two powerful and widely used modeling platforms, CC3D and Geant4, we delivered a novel tool that can give us the ability to simulate the dynamics of biological tissue in the presence of ionizing radiation, which provides a powerful framework for quantifying the biological consequences of radiation therapy. The developed tool has an advantage on that it has strong extensibility due to the exploitability of two modeling platforms. In this introductory methods paper, we described our modeling platform in detail and showed how it can be applied to study the application of radiotherapy to a vascularized tumor.

## 1. Introduction

Cancer is a significant health problem and is the leading cause of death associated with the aging of the population and lifestyle. Radiotherapy aims to sculpt the optimal isodose on the tumor volume while sparing healthy tissues. The benefits are threefold: patient cure, organ preservation, and cost-efficiency [1]. However, there are some challenges that pose to radiation oncologists as they attempt to predict and interpret the biological consequences of radiotherapy treatment. The mathematical modeling serves a suitable approach to overcome these difficulties since modeling can help us understand the underlying action mechanism of the biological effects of radiation. In terms of treatment planning in radiation therapy, modeling provides the means for optimizing radiation applications: the optimization methods implemented in treatment planning systems can be used to obtain the best achievable balance between the intended effects and inevitable side effects using the quantitative mathematical model. Radiation-induced biological effects result from complex mechanisms that involve a multitude of processes running at very different spatial and temporal scales and need a multiscale modeling approach, such as modeling of radiation effects on subcellular scales, modeling of cell killing, and modeling biological effects in tissues or organs (reviewed by Friedland et al [2]).

However, this is a challenging task because even using some rough mechanistic models of how radiation interacts with cells, such as the current LQ-centered approaches, it remains quite challenging to extrapolate this at tissue level, considering the tumor heterogeneity and microenvironment of tumor. The integrated stochastic model couples cell biology with radiation transport and has shown to quantify the biological consequences of radiation by taking into account the physical and biological processes of irradiated tumors at cellular level in a comprehensive way, which could be adopted to tackle the difficulties mentioned above.

Another aspect of modeling radiation-induced cellular effect needed to be mentioned is that modeling the proper spatiotemporal environment is crucial as a cell’s behavior depends strongly on its microenvironment. In other words, given the effects of radiation on a single cell, it is very hard to quantify the effects of these doses at the tissue scale. This is precisely a situation where multiscale quantitative tissue modeling can play an important role in designing better radiotherapy treatment protocols. Recently, Powathil et al. developed a multiscale mathematical model of chemotherapy treatment, incorporating cell-cycle mediated intracellular heterogeneity and external oxygen heterogeneity to study the effects of cell-cycle, phase-specific chemotherapy, and its combination with radiation therapy [3]. In their model, the radiation dose was directly assigned to cells according to the dose delivery scheme, ignoring the radiation transport calculation to obtain the individual cell dosage, resulting in a uniform dosage. However, experimental results indicate that there is a stochastic distribution of cell dose after irradiation [4]. Although, some researches considered the stochastic characteristic of cell dose [5][6], the models for quantifying the radiation interactions in cell is simply relying on a Poisson distribution. It is not accurate enough to describe the radiation dose delivering mechanisms of different types of radiation as the model neglects the difference between the radiation tracks by different types of radiation.

While there exist many radiation therapy models, very few offer the ability to study the impact of radiation on individual cell and yet, at the same time, give the ability to simulate entire tissue. CC3D + RADCELL/Geant4 offers such capability. Our computational platform is based on two key software components: CC3D and Geant4. CC3D focuses on cell-based tissue models (where each biological cell is modeled as an individual entity), while Geant4 is a well-validated Monte Carlo Simulation tool for studying the interaction of elementary particles (including photons) with matter. Merging the two tools into one modeling platform gives us an ability to simulate how dynamics of biological tissue is in the presence of ionizing radiation. At each step of the simulation, we assess the current effect of the radiation on simulated cells and adjust the properties of affected cells and their environment accordingly. Those adjustments have an effect on the temporal evolution of the cellular patterns. Thus, by having a way to simulate in detail how radiation affects individual cells, we can build more realistic, and hence more predictive models of radiotherapy.

This is a methods paper where we present a novel computational framework that we have developed with a few example applications. It is worth noting that this paper mainly is mainly focusing on its technical feasibility of implementing integrated simulation by combining RADCELL and CC3D. We acknowledge that there is still some improvement space for achieving a higher biological accuracy in simulation. For instance, in this work we don’t take care of the molecular level simulation, and we focus on cellular level. In the current design of our model, we don’t simulate the DNA damage repair and aberrant repair, etc. which is important in modeling radiation-induced DNA damages. Another thing is that we don’t simulate the radiation-induced DNA damages of mitochondria, and this may underestimate the total DNA damages of radiation to some cell lines [7][8]. But CC3D can conduct the sub-cellular system simulation [9]. If we have the pathway known, we can simulate the DNA damage and repair on the molecular scale. Due to the easy extensibility of RADCELL/CC3D, extra models can be added to address these issues. The primary focus of this paper is on the implementation of our platform, details of the modeling applications will be discussed in a subsequent paper.

In this paper, we firstly present how the simulation framework is developed, and then we devote the last sections to present how the developed simulation framework is used to simulate the cell and tissue response after irradiation, and specifically, an example of vascular tumor irradiation using microbeam is studied.

## 2. Methods and Material

In this section, we talk about how the coupled simulation platform is developed. The coupled simulation method includes radiation transport simulation, and cell biology simulation, to simulate the cell response after irradiation. The radiation transport simulation is implemented using RADCELL/Geant4, and cell biology simulation is implemented using CC3D. In the simulation, CC3D serves as a master control of the whole simulation process, and all the simulation process is controlled through the user interface of CC3D which supports Python scripting.

### 2.1 CompuCell3D

Simulations that focus on emulating cellular behavior can be divided into two broad categories: single-cell and multi-cell simulations. For single cell simulation, we implement detailed models of “intra-cellular dynamics” and typically this is accomplished by either using ordinary differential equations (reaction-kinetics models), reaction-diffusion equations (such as models implemented using Virtual Cell modeling platform http://www.vcell.org) or particle-based models (e.g., based on the MCell modeling platform http://mcell.org/) that are inspired by molecular dynamics models but are more coarse grained and operate at much larger time-scales.

While single cell-models are capable of explaining many phenomena observed at a single cell-level, the computational costs of running those models are very high, and consequently, building tissue models based on either of the approaches mentioned above are not practical. To deal with those computational limitations, multi-cell tissue models make a series of simplifying assumptions. The level of detail with which we represent a single cell in the multi-cell models usually determines the multi-cell model size (measured in terms of number of simulated cells). The simplest representation of single cell is a point in space (implemented in the Cellular Automata Model [10][11] or the Center Model [12]) allows representing relatively large number of cells in the simulations, while methods that aim to model cells as spatially extended domains capable of modeling cellular shapes and shape changes have to perform more computations. Yet, on modern computers, the more detailed multi-cell simulations such as those based on Cellular Potts Model (CPM) [13], or Subcellular Element Model [14] can model tissue fragments that are big enough for running fairly realistic virtual assays.

In our developed simulation platform, we use CC3D as our cell biology simulation platform, due to its easy implementation and powerful simulation capabilities. CC3D provides a user-friendly programming environment in order to develop simulation models. The fundamental structures and functions are developed using C++ for speed, but in practice bulk of CC3D simulation specification takes place in Python which makes the daunting task of constructing the model much simpler. CC3D also provides a dedicated Model Editor (Twedit++) that significantly reduces the burden of setting up the model by offering modelers with a set of code-assistants for the most common modeling tasks. In addition to simulating cellular mechanics, CC3D allows coupling of multiple modeling scales. It provides a set of PDE solvers that operate at tissue and organ scale while the reaction-kinetics solvers that CC3D provides (implemented using libroarrunner [15]) allow modeling of intracellular-phenomena. By appropriate linking the modeling scales (from molecular to organ-level), we can build very sophisticated models of tissues and thus gain insight into how disease processes in tissues originate and progress [16][17][18].

CC3D implements CPM [13] also known as Glazier-Graner-Hogeweg Model (GGH) [19][20][21][22]. In CPM, we represent generalized cells as spatially-extended domains that reside on a lattice. Each generalized cell in CC3D is made up of lattice sites, which are referred to as pixels or voxels, and can represent biological cells, compartments of cells or other biological objects such as the extracellular matrix. Each lattice site is represented by a vector of integers *i*. When a pixel is part of a generalized cell, the cell index is referred as σ(*i*), and the cell type is called *τ*(σ(*i*)), shown as in Figure 1. The description of interactions between generalized cells is implemented using the effective energy which determines many characteristics such as cell size, shape, motility, adhesion strength, and the reaction to gradients of chemotactic fields.

**Figure 1:**
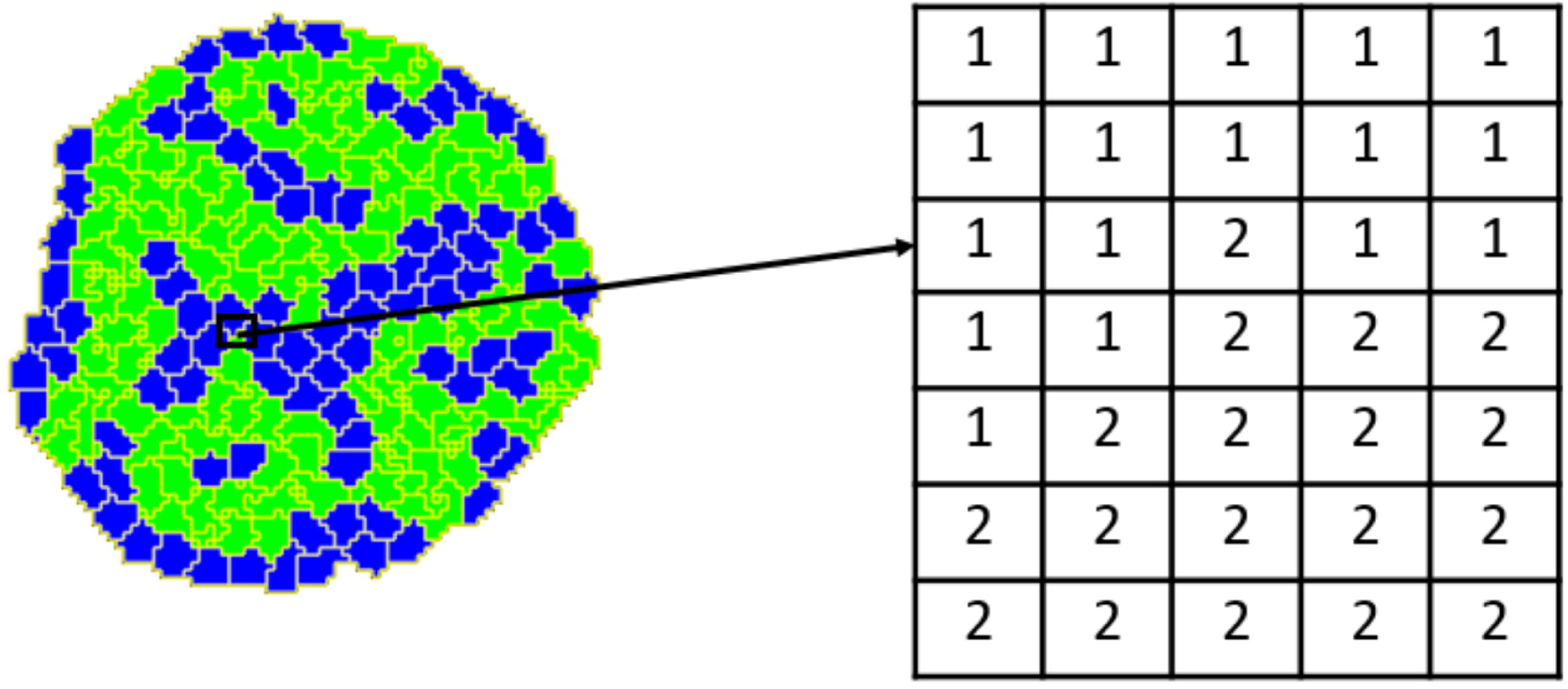
Detail of a typical two-dimensional CPM cell-lattice configuration. Each colored domain represents a single spatially-extended cell. The detail shows that each generalized cell is a set of cell-lattice sites, *i* with a unique index, *σ* (*i*) here 1 or 2. The color denotes the cell type, *τ* (*σ* (*i*)).

A simulation progresses by attempts of generalized cells to extend their boundaries in an effort to minimize the effective energy. These are called index-copy attempts because they try to change the cell index of a neighboring pixel to that of its cell type. The success of the index copy attempt is dependent upon a Boltzmann acceptance function which takes into account the change in energy, which is illustrated in Figure 2.

**Figure 2:**
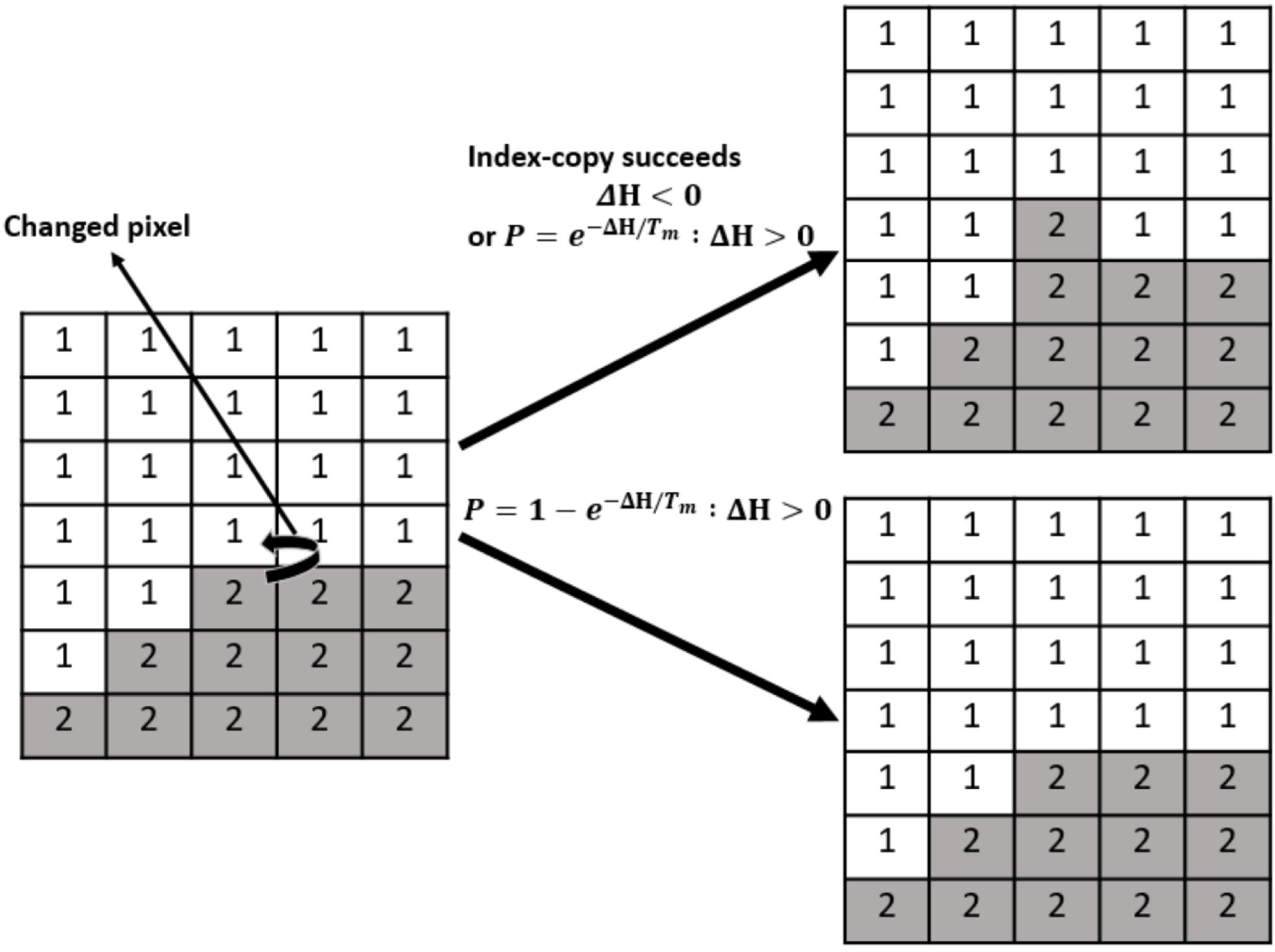
CPM representation of an index-copy attempt for two cells on a 2D square lattice-The “grey” pixel (source) attempts to replace the “white” pixel (target). A pixel, chosen at random, of type 2 cell makes a pixel-copy attempt to a neighboring pixel of a type 1 cell. The pixel-copy attempt is accepted and the neighboring pixel transitions from type 1 to type 2. If the attempt is not accepted, then there will be no pixel transitions.

The detailed description of the effective energy calculation and the simulation dynamics is referred to Appendix 1: CompuCell3D general introduction.

### 2.2 Geant4 in General

Geant4 is a freely available software for performing Monte Carlo simulations of the interactions of energetic particles in matter [23]. Geant is an acronym for “GEometry ANd Tracking”. The Geant4 is developed by CERN for the purpose of simulating the interaction of high energy particles in large accelerators. Since its development, Geant4 has continued to evolve and expand. The original toolkit implemented methods for handling the fundamental aspects of physical simulation: geometry, materials, particle interactions, particle generation, particle tracking, generation and storing of event and tracking data, and detector and tracking visualization. From these building blocks, the code has expanded widely. Its object-oriented nature allows the end-user to make use of the existing framework by instantiating useful class systems while maintaining a unique and personal application framework.

In the developed simulation platform, Geant4 serves as the radiation transport solver for our specific simulations, such as calculating the energy deposition points inside cells. The main reason for choosing Geant4 is that it is an open-source Monte Carlo simulation platform, and it provides many flexibilities for users, and also the low energy DNA damage simulation physics process, Geant4-DNA.

### 2.3 RADCELL

In order to couple Geant4 solvers with CC3D, we need a “bridging” code that extracts current geometry of the CC3D simulation (including specification of the individual cells) and ports it to a Geant4 solver. In other words, we need to “translate” representation of biological cell as implemented in CC3D into a computational representation of the cell that Geant4 can work with. This task is accomplished using our newly developed module, RADCELL. RADCELL is a radiation transport simulation module developed for conducting radiation transport simulation in cells. It can simulate the cell dose and cell DNA damages, such as single-strand breaks (SSBs) and double-strand breaks (DSBs). The RADCELL is developed based on the microdosimetry example of Geant4. The functionalities of RADCELL are referred to Appendix 2: RADCELL general introduction.

The primary function of RADCELL is to calculate the radiation dose to cell organelles, and DNA damages to cells. In this work, we propose a three-dimensional (3D) cellular compartment model, which incorporates two cellular compartments including nucleus and cytoplasm. We use the sphere to approximate the cell shape which substantially simplifies the complexity of cell geometry but there is no significant big accuracy penalty [24]. The nucleus is modeled as a sphere which is located at the center of the cell. The size of the cell and the nucleus can be customized according to the biological cell which will be studied during the simulation.

During the radiation transport simulation, the energy deposition information in each cell will be collected, and that information could be used to quantify the cell dose and DNA damages. For a better illustration, here we list the information of one energy deposition point as below:

- eventID, which is used to indicate in which event (equivalent as one irradiating particle during cell irradiation) in which the energy deposition interaction happens
- cellID, which is used to indicate in which cell energy deposition point locates
- edep, which is used to indicate the energy deposited at the energy deposition point
- pX, which is used to indicate the *x* coordinate of the energy deposition point
- pY, which is used to indicate the *y* coordinate of the energy deposition point
- pZ, which is used to indicate the *z* coordinate of the energy deposition point
- affectedCellOrganelle, it is used to indicate in which cell organelle the energy deposition point locates.

#### 2.3.1 Cell Dose Tally and DNA Damage Tally

After we obtain the energy deposition information in each event, then we process these data to obtain the cell dose tally and cell DNA damage tally. In RADCELL, cell dose tally is quantified by summing the total energy deposited in cell and dividing the energy deposited by cell mass. For cell DNA damage tally, the SSB and DSB yields are quantified using a clustering algorithm, i.e., DBSCAN (Density-based spatial clustering of applications with noise) [25] for processing the energy deposition information inside nucleus. The clustering algorithm is a popular method for quantifying DNA damages yield, which is discussed in [26][27][28]. A detailed description of the development of the algorithm for cell dose tally and DNA damage tally based on Geant4 simulation is discussed in our previous work [29][30]. Here we give an example of quantifying the cell dose and DSB yield of a multicellular spheroid under the irradiation of monoenergetic electron source. The irradiation setting is shown in Figure 3a, which is performed using a parallel plane electron source. Figure 3b suggests the simulated cell dose distribution of the spheroid after the irradiation. Each cell is rendered as a spherical ball, and the color bar is used to scale the cell dose quantity. We can examine the heterogeneity of the cell dose distribution within the spheroid system, which shows that the dose of cells of spheroid under uniform irradiation is not always uniform. Figure 3c indicates the simulated cell DSB number distribution of the spheroid after the irradiation. Similarly, the color bar is used to scale the DSB number quantity. We also can observe that the DSB number distribution is not uniform across all cells.

**Figure 3:**
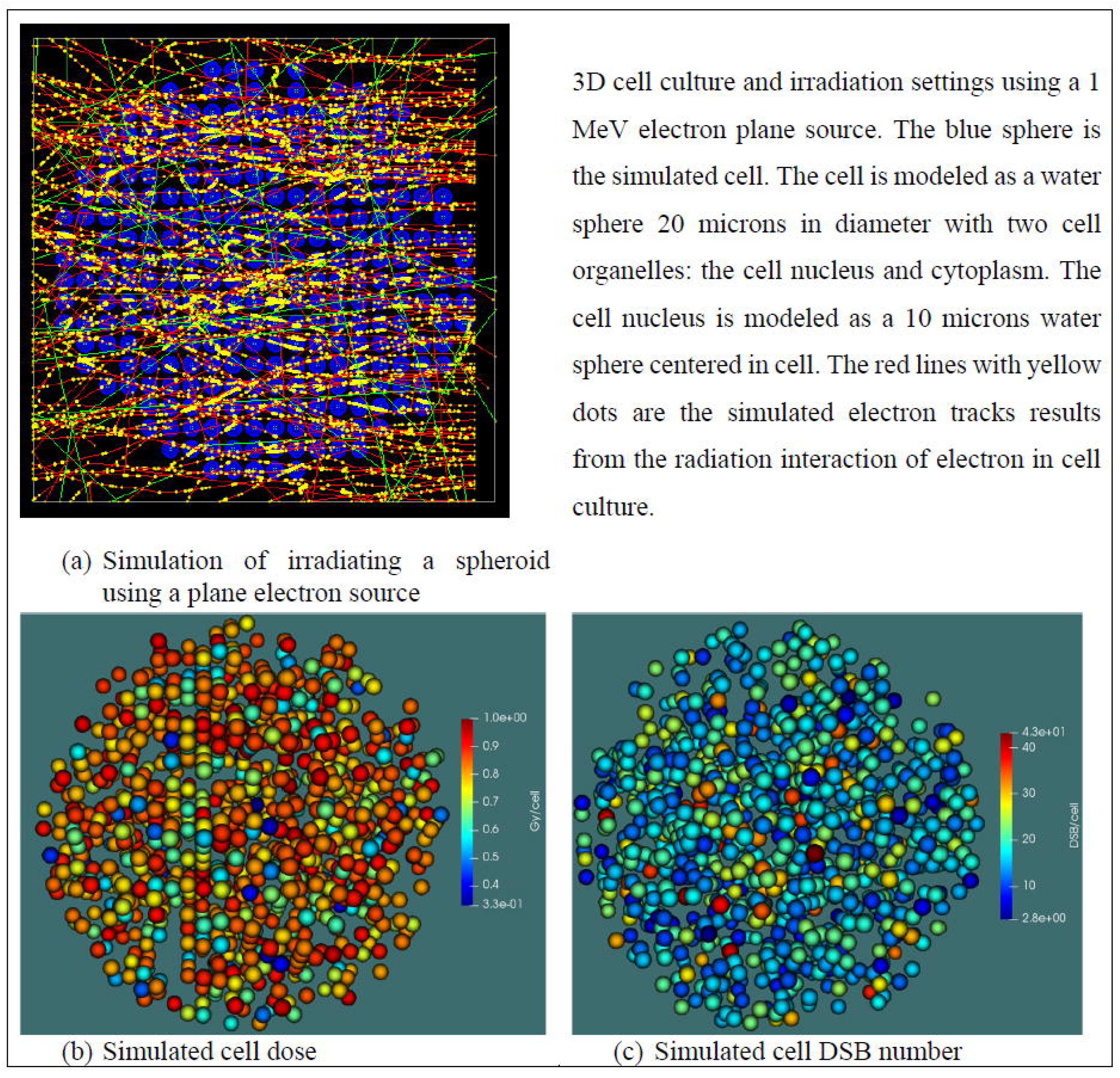
Simulated cell dose and DSB number distribution of a spheroid with 1000 cells after irradiation. The total history number in the simulation is 1,000,000, and the shape of the plane source is set up as a square shape and its size is set up as bigger than the size of cell culture (Figure was made from our previous publication [29] with permission).

In the current functionality for modeling radiation-induced DNA damage in RADCELL, there are a few limitations needed to be mentioned. For the DNA damage repair of cell, we don’t simulate the DNA damage repair and aberrant repair, etc. Modeling approaches based on biochemical kinetic equations have been proposed for representing the course of the base and nucleotide excision repair systems, which is used by cells to remove base damages and strand breaks and more bulky lesions, respectively [2]. The DSB damage repair mechanism, such as nonhomologous end joining (NHEJ) could be modeled to simulate the DSB repair in cell [31]. We don’t simulate the radiation-induced DNA damages of mitochondria, and this may underestimate the total DNA damages of radiation to some cell lines [7][8]. Due to the easy extensibility of RADCELL/Geant4, extra models can be added to address these issues.

#### 2.3.2 Cell State Transition Model

In this work, we use a cell state model to quantify the temporal transition of the possible cell phenotypes after irradiation [32][33][34]. We defined three major cell states: ***Healthy, Arrested***, and ***Dead*** [30]. A ***Healthy*** cell maintains its basic functionality or keeps a proliferative state with no or very light damage. An ***Arrested*** cell has its cycle halted in a specific cell-cycle phase. A ***Dead*** cell has suffered irreparable damage and suspends material exchange with extracellular matrix (ECM). Each of these cell states differs depending on the cell’s phase in the cell cycle: G1, S, G2, or M. The allowed state transitions are:

- From Healthy to Arrested or Dead.
- From Arrested to Dead or Healthy.

Dead cells stay Dead. Transitions depend only on a cell’s current state.

These state transition rules are based on radiation biology experiments. For instance, 1) A high dose causing direct cell death corresponds to the state transition from Healthy to Dead; 2) A moderate dose causing cell-cycle arrest corresponds to the state transition from Healthy to Arrested; 3) Cell apoptosis after failed cell damage repair during cell-cycle arrest, corresponds to the state transition from Arrested to Dead; 4) A dead cell does not have capacity to repair damage, so the Dead state is persistent. The mathematical description of the cell state transition model is referred to Appendix 3: Cell State Transition Model.

During simulation, the cell state transition is taken as a continuous stochastic process evolving with time after cell irradiation. Firstly, the external perturbation energy Δ*E* for each cell is updated, then the probability of corresponding cell state transition is calculated based on Δ*E* in the time step. Secondly, the cell state transition decision is made based on the calculated transition probability according to rejection sampling rule [35]. An example of updating the cell state in one time step is shown in Figure 4. Basically, the cell phase transition and cell state transition of cell after irradiation are followed and updated using a cellular automaton method.

**Figure 4:**
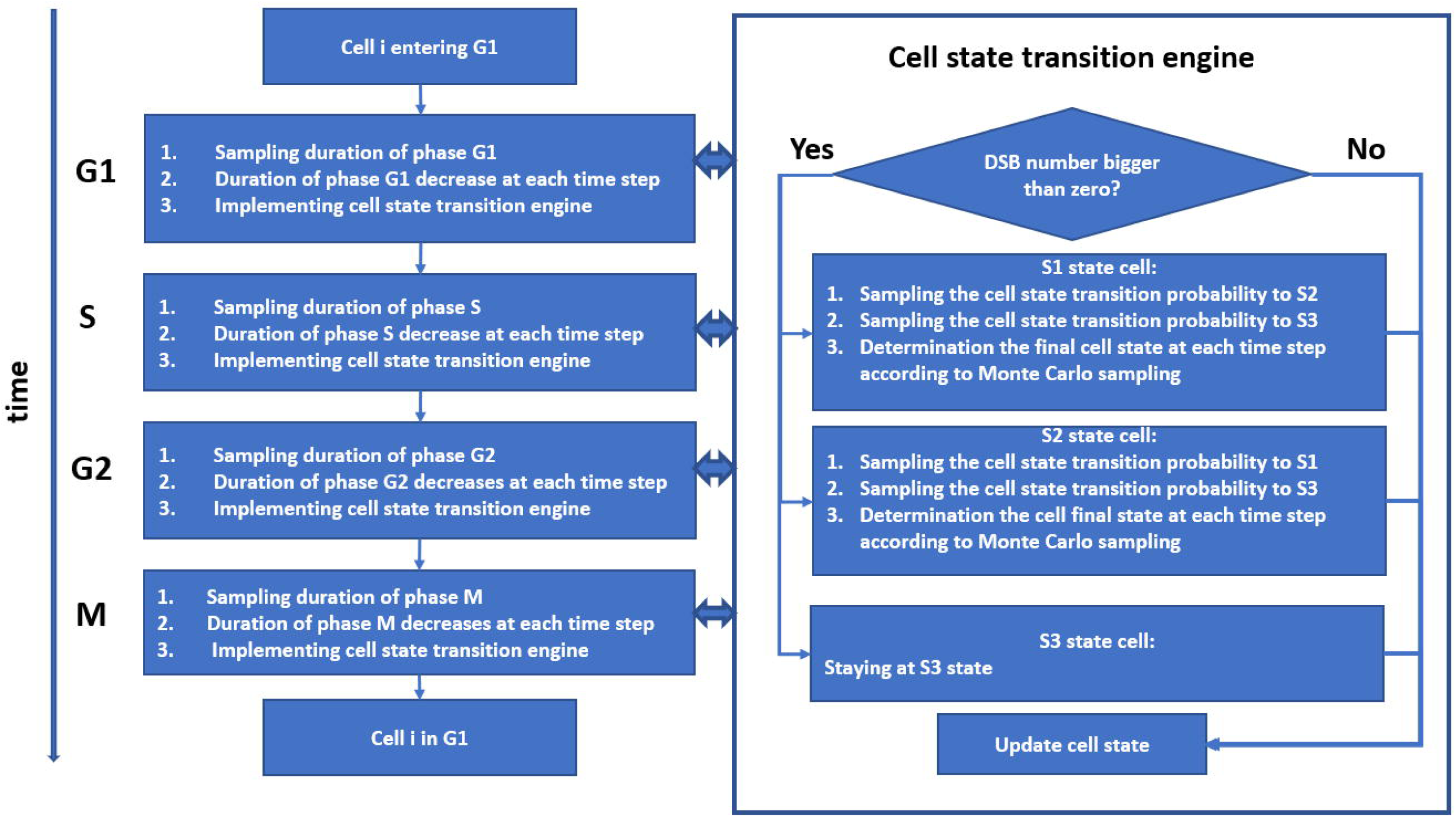
Process of updating cell state in one time step. During simulation, the state of all the cells will be updated in each time step which could be defined in the simulation. Cell phase transition is modeled by comparing the sampled cell phase duration and time of cell staying in the cell phase. If the time of cell staying is larger than the sampled cell phase duration, then cell phase transition will occur. If mitosis occurs, then two new cells will be added to the cell system. Cell state transition in each cell phase are also modeled through the defined cell state transition rules. In each time step, the cell phase and cell state transition are updated.

#### 2.3.3 RADCELL Implementation

The block-diagram of RADCELL integrating with Geant4 is shown in Figure 5. We first instantiate RADCellSimulation object and this object’s methods to build the simulation geometry and run the transport simulation. The RADCellSimulation supports two run modes: *gui* mode and *detailed* mode. When *gui* mode is chosen, RADCellSimulation will only run radiation transport, and the information of energy deposition points will not be collected and processed during the simulation. Typically, we use this mode to check whether the cell geometry is right or not. When a *detailed* mode is chosen, the RADCellSimulation will collect and process the information of energy deposition points during the simulation. The *detailed* mode should be chosen if we want to get the cell dose and DNA damage tallies.

**Figure 5:**
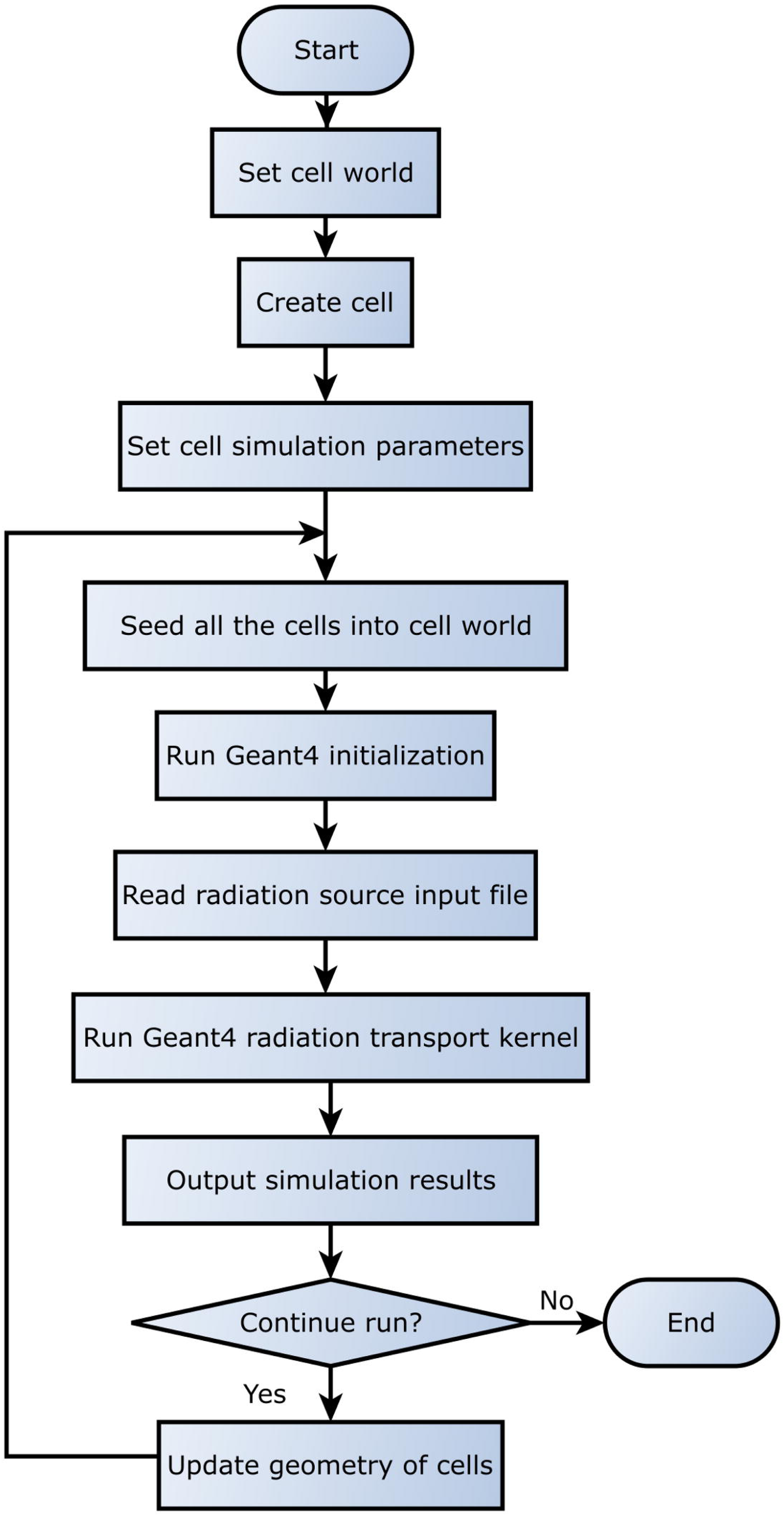
The block diagram of radiation transport simulation using RADCELL and Geant4. For radiation transport simulation of multicellular system, firstly, the computerized cells are created in the model, then conducting radiation transport calculation to obtain the simulation results, such as cellular dose and double-strand breaks. It is worth noting that the geometry in Geant4 should be updated if the multicellular system changed due to mitosis or cell death.

### 2.4 Coupled Simulation Using RADCELL and CompuCell3D

To enable seamless integration of RADCELL, Geant4 and CC3D, we have developed Python modules (using SWIG [36]) that allow using of all RADCELL functionality from the Python level, thus making it very easy to integrate with CC3D. Therefore, we can quantify cell dose and cell DNA damages and use those quantities to determine the cell state transitions for CC3D cells. This is significant, because using our approach, we can have the most up-to-date information about cell irradiation and thus can simulate with great level of detail how radiation impacts cell cycle or any other cell properties that can potentially be affected by increased levels of radiation. The block diagram of coupling RADCELL and CC3D is shown in Figure 6.

**Figure 6:**
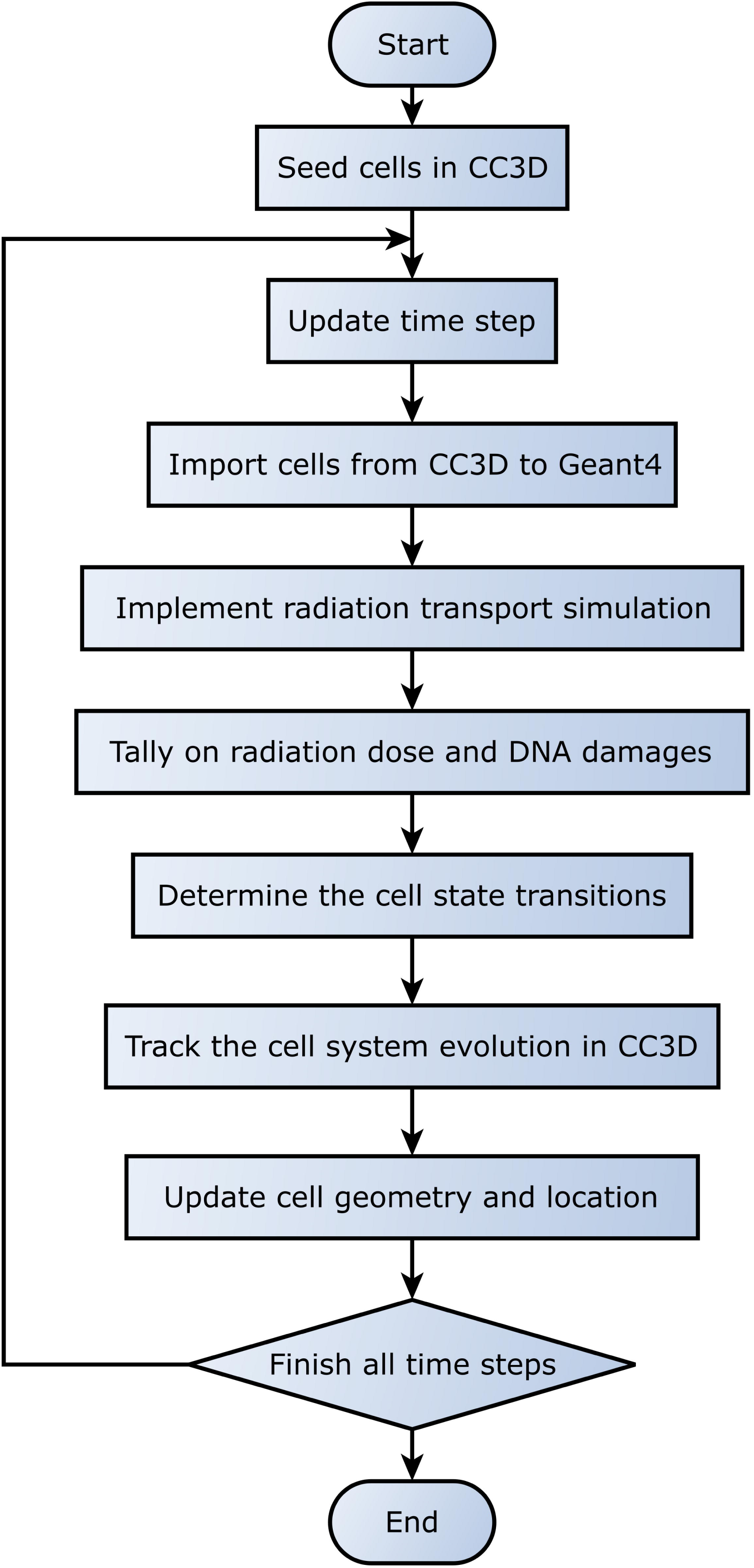
The flowchart of cross platform simulation using RADCELL/Geant4 and CompuCell3D. The cell position information is extracted from CC3D in order to build geometry of cells in Geant4. The CC3D conducts cellular system simulation based on the simulation results from RADCELL.

#### 2.4.1 Importing Cells into Geant4 from CC3D

Any attempt to couple separate simulators (in this case, Geant4 and CC3D) requires development of systematic methods that allow transferring information about simulated objects between the separate simulators. In our case, the challenge is how to represent CC3D cells in Geant4. To do that, we first extract CC3D cell position information and pass this information to Geant4. We then create Geant4 equivalents of CC3D cells in such a way that relative cell distances are preserved in Geant4. Getting correct positions of cells is crucial if we want to obtain the accurate radiation dose distribution for all the cells.

In Geant4, the cell has the same size as the biological cell for modeling the physical interactions accurately. The cells in Geant4 are seeded according to cells’ physical sizes. In this iteration of our software, we make a simplifying assumption about Geant4 cell shape and treat all the cells as spheres. After obtaining the cell position and size information, the cells are seeded into Geant4 for radiation transport simulation.

One of the potential problems is that the cells in Geant4 may overlap with each other if we simply seed the cells according to the center of mass (COM) of cells in CC3D. To overcome this problem, we keep the size of Geant4 cell constant and rescale the whole geometry to the point where all Geant4 spheres representing CC3D cells are non-overlapping. This rescaling has negligible effect of assessment of radiation damage. One way to think about it is to imagine that after this rescaling, the Geant4 spheres represent the entire CC3D cell but with a somewhat smaller CC3D cell volume (one that includes cell’s nucleus) that is directly susceptible to radiation effects.

Another thing to consider is the cell motion. Since CC3D is a Monte Carlo simulation technique, in every Monte Carlo Step (MCS) a given cell will appear in slightly different position. However, physically observed cell displacement are those that are observed every several MCS (the exact number of MCS interval depends on the simulation parameters [37]). We thus set a predefined MCS interval and use it to synchronize CC3D and Geant4 tissue layouts.

#### 2.4.2 RADCELL, CC3D, and Geant4 interoperability

After the radiation transport simulation (using Geant4), the cell dose information and cell DNA damage information are written to CSV files respectively. Then CC3D will read the dose and DNA damage information when they are needed for determining the cell state transition after irradiation. In the next releases of our software, we will eliminate the need to carry those CSV files and exchange information between Geant4 and CC3D using a more elegant solution.

When we run the simulation, CC3D is the master module that controls the execution of RADCELL and Geant4, as shown in Figure 7. The CC3D simulation (all internal modules of CC3D) is implemented in one operating system process, while the radiation transport simulation (using Geant4) uses another process which is different from the CC3D’s process.

**Figure 7:**
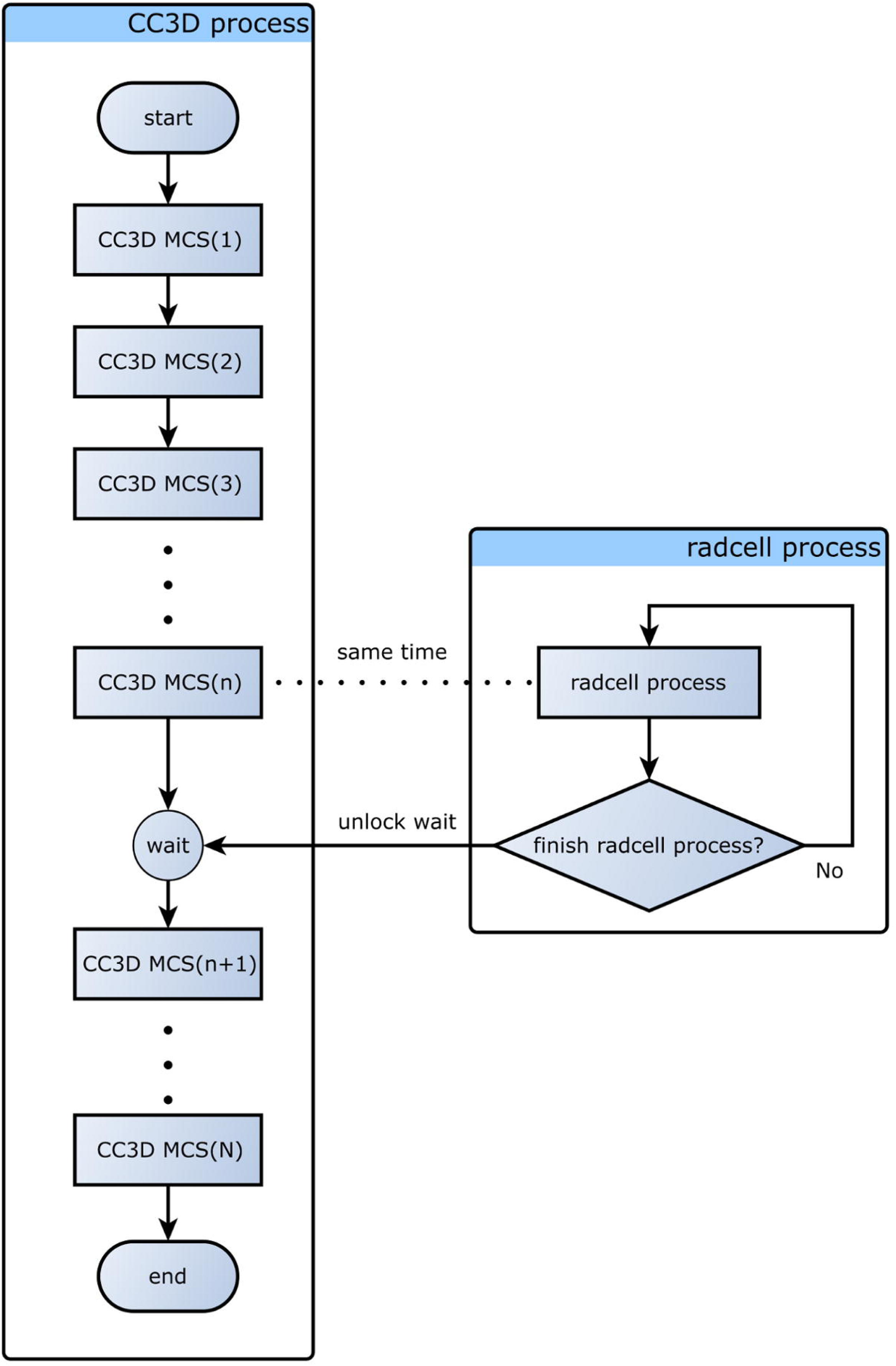
The coupled simulation process between RADCELL and CC3D.

As shown in Figure 7, at n^th^ MCS, in CC3D, RADCELL is launched to run the radiation transport simulation, so the radiation transport simulation results are just for the cells at n^th^ MCS. It is essential to ensure that while RADCELL (Geant4) executes radiation transport calculations, CC3D waits for the radiation transport results before proceeding further. Once CC3D gets radiation information, it will modify cell properties accordingly and proceed with the next n+1 MCS, and the whole process of interleaved CC3D+ RADCELL execution will continue to repeat until CC3D simulation is over.

## 3 Example Model

In this section, we talk about how to use the developed simulation platform to simulate the cell and tissue response after irradiation. We developed a simplified model of microbeam irradiation of a vascularized tumor. This model is provided as a demonstration of the capabilities of CC3D-RADCELL-Geant4 modeling platform. In this technical paper, we will not, however, explore those models in detail. This will be done in follow-up papers.

The typical Microbeam Radiation Therapy (MRT) treatments deliver microscopically discrete spatial dose distributions: 10-100 microns wide parallel beams with a separation of several hundred microns between adjacent beams. Because high-dose, high-precision MRT’s have potential to significantly reduce probabilities of healthy tissue complications [38][39], they are considered as a promising treatment concept. However, one of the significant challenges is the lack of comprehensive understanding of the underlying radiobiological mechanism. It is commonly acknowledged that MRT is based on the principle that healthy tissues can tolerate high doses of radiation in small volumes and that MRT can damage the blood vessels, cut off the tumor nutrient supply, and as a result, cause the tumor to die [40][41].

To better understand the radiological mechanism of MRT, we focus on two core areas: 1) understanding how radiation transport process of microbeam radiation affects tumor and healthy tissues, 2) understanding the tumor response after microbeam irradiation.

While microbeam radiation transport has been already addressed in several publications, e.g., [42][43], there has been somewhat less emphasis on how tumor behaves subject to MRT. Because the RADCELL-CC3D platform facilitates coupled tissue-radiation simulations, it is an ideal tool to conduct these sorts of studies.

In the remainder of this section, we present a detailed guide demonstrating all steps needed to build a model of radiation treatment of vascularized tumor. Our model simulates essential cell behaviors, microenvironmental components, and their interactions as well as radiation response after irradiation. The basic modeling methods and procedures are discussed as follows. The work is based on a simulation work by Swat et al. [22]. Some sub-models simulated in the example simulation are displayed in Table 1. It is worth noting that these models are just selected for demonstrating the capability of coupled simulation of our developed simulation framework. Due to the easy extensibility, other models could be easily added for achieving more functionalities.

**Table 1:** Models simulated in simulating the MRT treatment of vascular tumor

| Sub-models |  | Functionality | Platform used |
| --- | --- | --- | --- |
| Cell biology modeling | Multi-cellular model | Building tumor cells | CC3D |
|  | Reaction-diffusion model | VEGF signaling process; glucose reaction-diffusion process | CC3D |
|  | Cellular Potts Model | Cell-cell adhesion; cell volume constraint; cell surface constraints | CC3D |
|  | Michaelis-Menten expression from enzyme kinetic model | Cell growth and mitosis | CC3D |
| Radiation transport modeling | Radiation transport model | Parallel electron microbeam transport in tissue | RADCELL/Geant4 |

### 3.1 Vascular Tumor Model in CC3D

Firstly, we use CC3D to build the vascular tumor model. An initial condition for our vascularized tumor model is a cluster of proliferating tumor cells and a simple network of pre-existing normal vasculature. Initially, tumor cells proliferate as they take up diffusing glucose which the pre-existing vasculature supplies. In this work, we expect that the tumor cells (both in the initial cluster and later) are usually hypoxic and they can secrete a long-diffusion isoform of VEGF-A which is denoted as L-VEGF here. When the concentration of glucose drops below a threshold, tumor cells end up necrotic, step by step shrink, and ultimately disappear. A few preselected neovascular endothelial cells in the pre-existing vasculature respond both via chemotaxing toward greater concentration of proangiogenic factors and form new blood vessels by means of neoangiogenesis. The preliminary tumor cluster grows and reaches a maximum diameter characteristic of an avascular tumor spheroid. When the tumor grows to a certain size, we simulate the application of microbeam irradiations that kills certain fraction of tumor and vascular cells. We simulated deliveries of different irradiation schemes and implemented basic mechanisms that simulate tumor response and evolution after irradiation.

#### 3.1.1 Cell and ECM Types

Our model of solid tumor includes two main classes of generalized cells: tumor cells and stromal tissue. In this work, we stipulate four cell types: P: proliferating tumor cells; N: necrotic cells; EC: endothelial cells; NV: neovascular endothelial cells and ECM: an aggregate of stromal cells and ECM. This model focuses on small, early-stage vascular tumors rather on developed primary tumors, to identify the pattern of cell-behavior selection. While the average number of cells is far less than a real vascular tumor at any time, we can map these simplified model tumors onto real tumors either by considering each model cell to represent an ensemble of hundreds or thousands of real cells or by considering each model tumor to represent a peripheral microportion of a much larger tumor mass. In CC3D, we set the cell and field lattice dimensions to 50×50×80, the membrane fluctuation amplitude to 20, the pixel-copy range to 3, the number of MCS to 17,000, and choose UniformInitializer to produce the initial tumor and vascular cells, since it automatically creates a mixture of cell types.

#### 3.1.2 Chemical Fields

In this study, we consider two types of chemical fields, i.e., glucose, and VEGF-mediated signaling factors. The cluster of tumor cells form a spheroid, the nutrient and waste diffusion will limit the diameter of such avascular tumor spheroids to about 1 mm. The central region of the developing spheroid will become necrotic, with a surrounding layer to cells whose hypoxia triggers VEGF-mediated signaling events that initiate tumor neovascularization by way of promoting growth extension or nearby blood vessels [22]. We add a set of finite-element links between the EC cells to model the strong junctions that form between EC cells and NV cells chemotax up gradients of two diffusing isoforms of VEGF-A, i.e., S-VEGF and L-VEGF. Both EC cells and NV cells chemotax up S-VEGF, but only NV cells chemotax up gradients of L-VEGF. We assume that glucose includes a diffusing field representing glucose. The L-VEGF and S-VEGF include diffusion fields representing L-VEGF and S-VEGF respectively.

#### 3.1.3 Cell Interactions

Following basic assumptions of the CPM model [13], we simulate cell interactions by specifying effective energies associated with particular cell behaviors. The detailed description of cell interactions is referred to Appendix 4: Cell Models Used in CC3D Modeling.

#### 3.1.4 Cell Growth and Mitosis

In this study, we assume that glucose is the main growth-limiting substance for tumor cell. The L-VEGF is the main growth-limiting substance for neovascular cell. The concentration rate of glucose could be described by the reaction-diffusion equation. Cell growth and contact-inhibition growth of NV cells are modeled in this work. The detailed description of the models could be referred in Appendix 4: Cell Models Used in CC3D Modeling.

### 3.2 Radiation Transport Using RADCELL

In our coupled simulation, we import RADCELL directly into CC3D as a Python module. A snippet of code is shown in Figure 8. The few lines of codes importing the RADCELL package into CC3D. Then we define a CC3D steppable, RadiationTransport, to do the radiation transport simulation for cells. The functionality of CC3D steppable class is fully described in [21].

**Figure 8.**
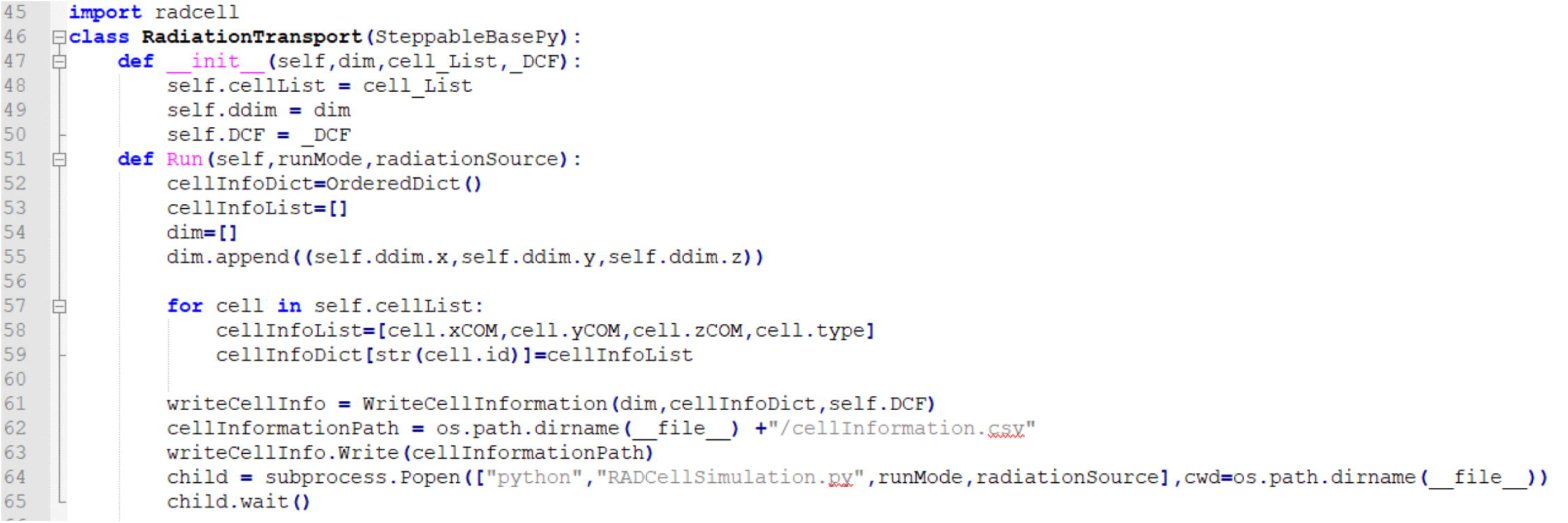
Sample CC3D steppable class conducting radiation transport simulation using RADCELL. The radiation simulation is interleaved with the simulation of cellular pattern evolution. After each application of radiation in our model, we query the RADCELL module to provide information about cell dose and DNA damages for every biological cell present in our models.

We use cell dose and cell DNA damage information to determine the cell state transition in CC3D which typically amounts to altering cell parameters such as cell type, cell target volume, and cell target surface. In general, every parameter that describes cellular behavior can be altered, and it is up to a modeler to come up with a reasonable way of doing it.

The value of combining radiation transport simulation within CC3D is that we can obtain the most up-to-date information about radiation damage and consequently build a more realistic radiation treatment model.

#### 3.2.1 Radiation Source

To model the MRT irradiation, we chose a square plane with 50 µm width as a radiation source. This dimension is in line with the width of the parallel microbeam used in MRT. Typical energies of X-ray radiation used in MRT range from 50 keV to 600 keV. Considering that the dose of X-ray is induced by the secondary electrons, we make a simplifying assumption and use 600 keV electron to model the 600 keV X-ray. It is worth noting that this is a simplification, but it should give us a good “first-order approximation” of the real phenomena.

#### 3.2.2 Importing Cells into Geant4 from CC3D

During the simulation of radiation transport, we extract the cell position in CC3D and pass this information to Geant4 to create proxies of CC3D cells in Geant4 simulator. Since we preserve the geometry of the CC3D tissue, the information about radiation damage we get from RADCELL/Geant4 should be accurate enough that we can assess radiation effects on each cell.

### 3.3 Cell State Transition

In our simplified model, we consider two cell states, i.e., healthy state (*S*_1_) and dead state (*S*_3_). We define the cell state transition rule according to two types of conditions of cell, i.e., microenvironmental factors and irradiation. Experimentally, microenvironmental factors, including mechanical stress, hydrostatic pressure, low pH, and starvation, can cause temporary or permanent changes in tumor cells. In this model, however, we only include cell state transitions due to nutrient availability and radiation dose. During a period of starvation (in a low-nutrient regime), if the glucose concentration is lower than the threshold, then tumor cell dies. For nutrient availability condition, we simply calculate the glucose concentration for tumor cells at each MCS and compare the concentration with the threshold concentration. For quantifying the cell state transition after irradiation, we only consider the direct effect of irradiation. The cell state transition rule for glucose concentration and irradiation is referred to Appendix 5: Cell State Transition.

### 3.4 Simulation Parameter

CPM simulations measure simulation time in terms of MCS, and the conversion between MCS and experimental time depends on the average cell motility. Biologically, MCSs are proportional to the experimental time [17]. We can relate the simulation’s MCS time-scale to minutes by comparing cell-migration speeds in simulation to typical cell-migration speeds in experiments. In this study, we use a 3D voxel with a side of 4 *μ*m, so the tumor cell volume is 64 *μ*m^3^. Since the experimental human umbilical vein endothelial cells (HUVEC) speed is about 0.4 *μ*m /min [22], and the cells in this simulation move at an average speed of 0.1 pixel/MCS, so one MCS represents 1 minute. Here, by knowing the time scale conversion between the CC3D time scale and the experimental time scale, we use MCS as the time unit for all the parameters related to time hereinafter in this work. The simulation parameters are referred to Appendix 6: Simulation Parameters.

### 3.5 Simulation Results

#### 3.5.1 Tumor growth Without Irradiation

We run otherwise identical simulations with and without MRT irradiation to study how MRT irradiation affects tumor growth and morphology. To begin with, we run the simulation to see how the tumor system will evolve without irradiation. The simulation results serve as a control for evaluating the irradiation effectiveness of MRT.

As shown in Figure 9b, we initialize the tumor cell cluster and two crossing vascular cords. We also add two NV cells to each vascular cord, 25 pixels apart. Without irradiation, the tumor system follows its typical biological evolution path, and we can see that the tumor grows bigger and bigger with the glucose update. We evaluate tumor growth by analyzing the number of proliferating tumor cells with respect to time. The simulation of tumor growth without irradiation showed an apparent increase of proliferating tumor cells with time, as shown in Figure 9a. Though initially we only seed two NV cells, we can observe that the neovascular cells are undergoing proliferation with the simulation time going on, and this is triggered by VEGF-mediated signaling events that initiate tumor neovascularization by promoting growth and extension (neoangiogenesis) of nearby blood vessels. We also can observe that some tumor cells die due to lack of nutrients, which is in line with the biological observations in experiments, that the nutrient diffusion limits the diameter of the cluster of tumor cells, and the central region of the growing spheroid becomes necrotic [22].

**Figure 9:**
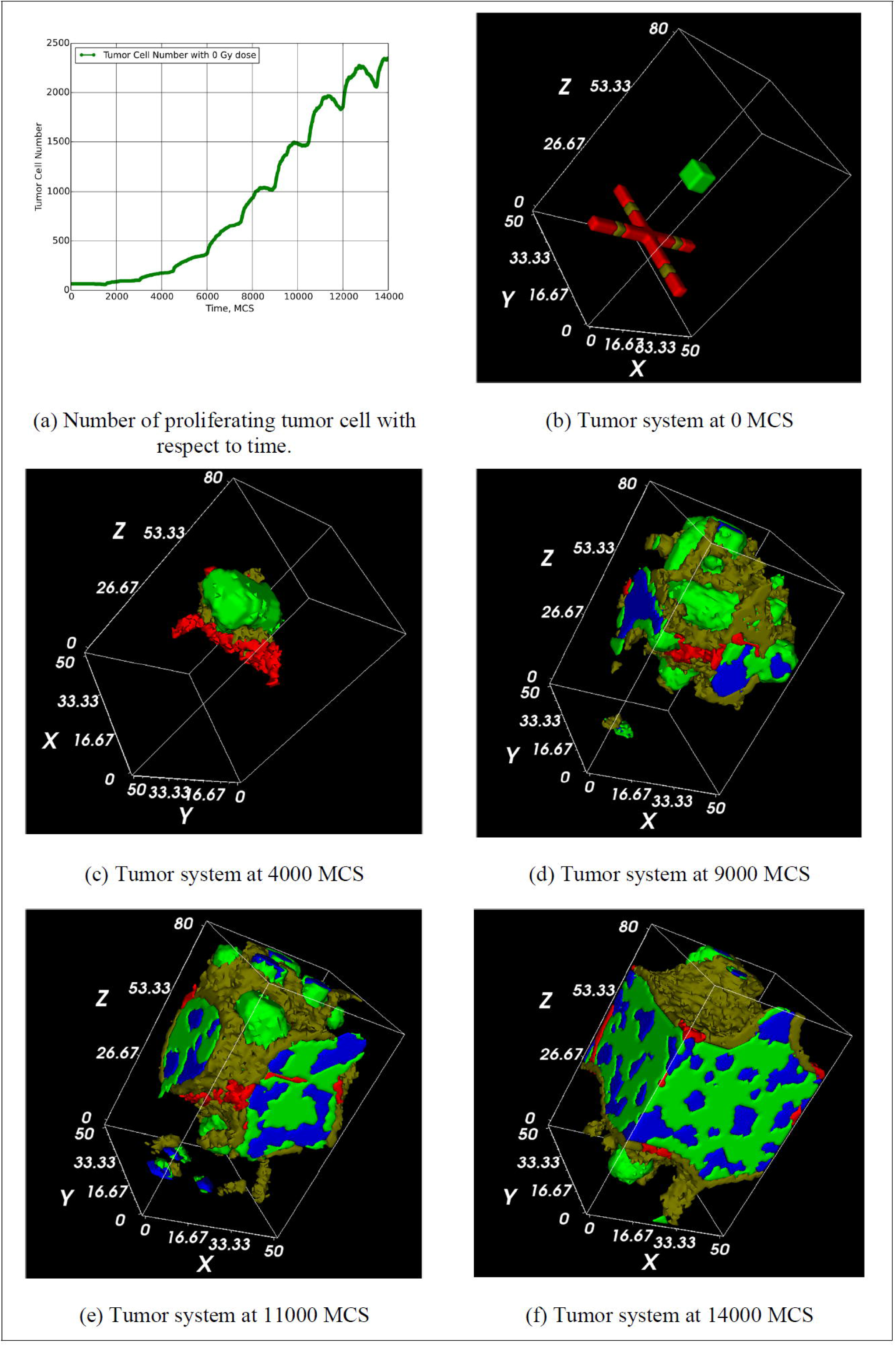
Two-dimensional snapshots of the vascular tumor simulation. Red: Vascular cells, grey: neovascular cells, blue: necrotic cells. The spatial unit used in CC3D is pixel, and one pixel equals 4 µm in this work. (a) shows the tumor growth curve with respect to tumor evolution time which has MCS as its unit. (b) shows tumor system at 0 MCS (starting time) rendering in CC3D visualization system. (c), (d), (e), and (f) show the tumor system at 4000, 9000, 11000, and 14000 MCS, respectively.

#### 3.5.2 Tumor Growth with Irradiation

For radiation therapy, radiation could be considered as an external perturbation agent to the tumor system. As we just simulate, without irradiation, the tumor system follows an unperturbed biological growing path, which indicates that the CC3D model can capture the basic biological characteristics of vascular tumor growth. Knowing this, we are more interested in knowing how the vascular tumor respond to the irradiation. Here, we conduct a simulation for simulating the tumor growth under different irradiation schemes.

In this simulation, we use a single planar microbeam source to irradiate the vascular tumor from an early time (starting from 1000 MCS), and the dose is delivered in five fractions equally. The total dose is from 5 Gy to 30 Gy. The dose delivering time is at: 1000 MCS, 5000 MCS, 7000 MCS, 9000 MCS, and 11000 MCS.

The vascular tumor at 1000 MCS is a relatively small cluster, as shown in Figure 10a. During simulation, when a vascular tumor evolves to this time, CC3D automatically launches the radiation transport simulation by calling the functions in the RADCELL module. The tumor geometry is extracted from CC3D and imported to Geant4 for radiation transport simulation, as shown in Figure 10b. It is worth noting that the materials of all the cells and the materials between cells are modeled as water in RADCELL. The radiation track structure of the single planar microbeam irradiation is shown in Figure 11.

**Figure 10:**
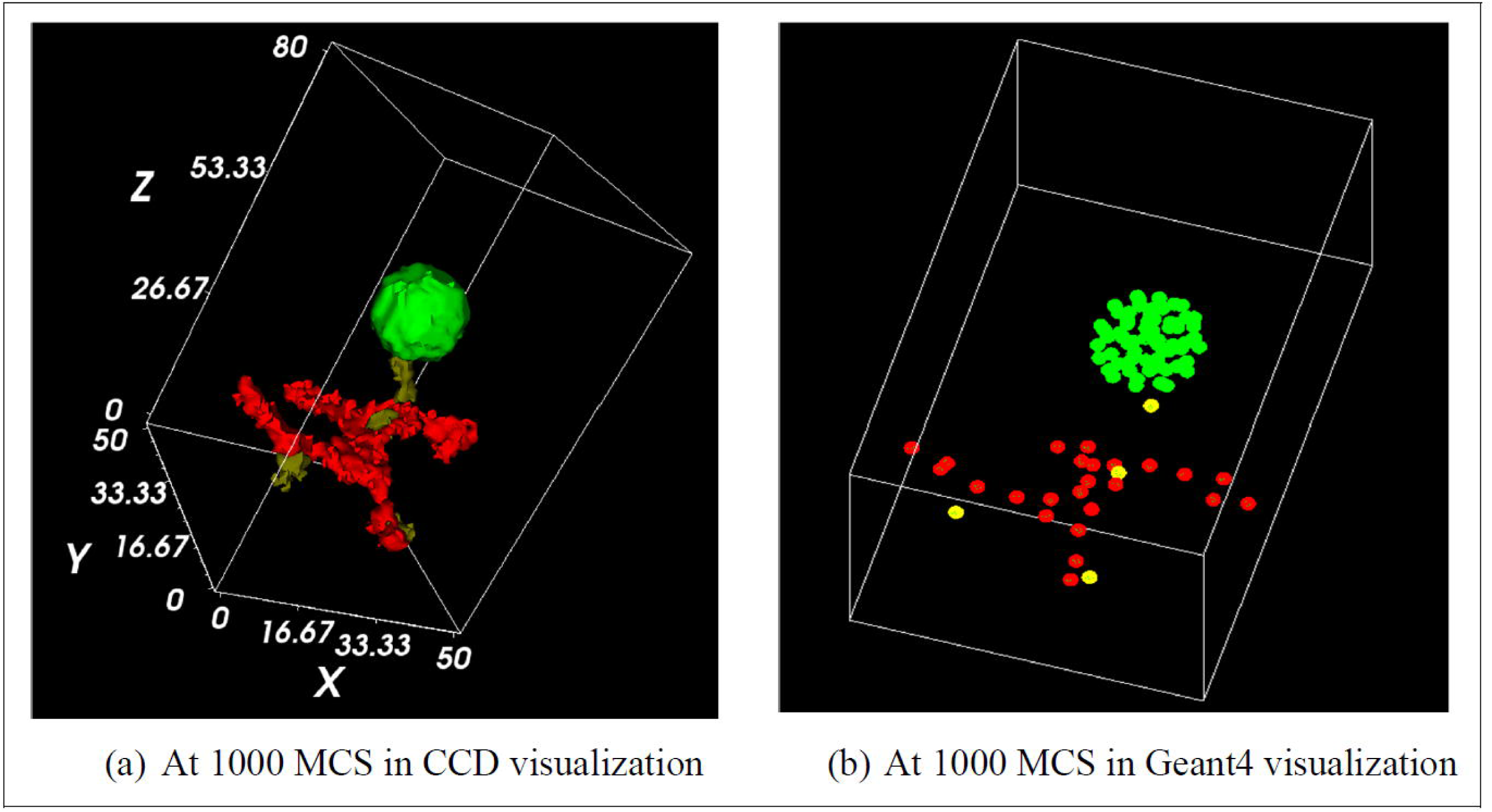
Two-dimensional snapshots of the vascular tumor simulation. The spatial unit used in CC3D is pixel, and one pixel equals 4 µm in this work. (a) is the tumor visualized in CC3D, and (b) is the tumor visualized in Geant4. Red: vascular cells, grey: neovascular cells, blue: necrotic cells.

**Figure 11:**
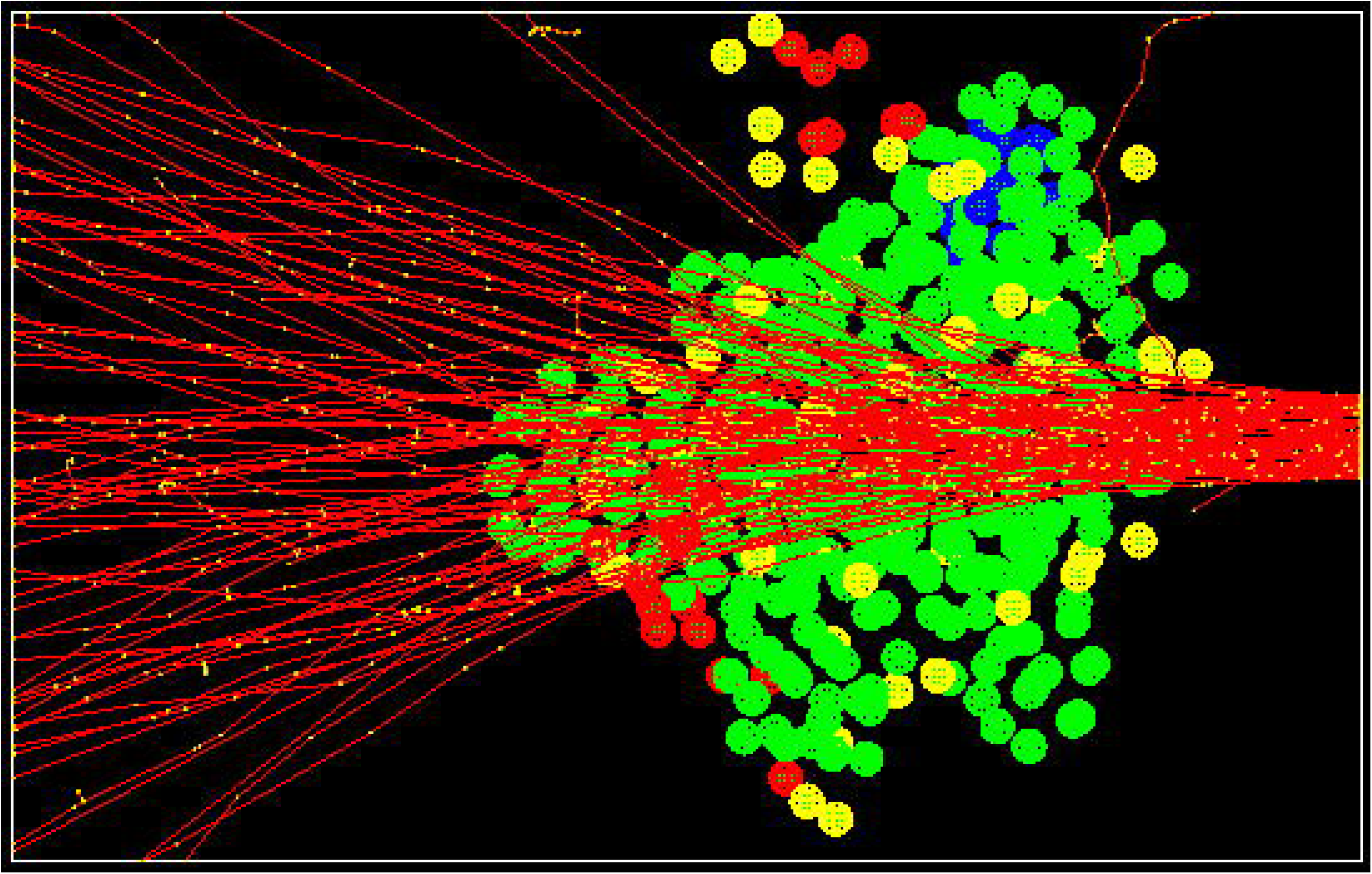
Single planar MRT irradiation on vascular tumor. In Geant4 platform, the red lines are the tracks generated by electrons, and the yellow points are the energy deposition points after radiation interactions.

Each time after radiation transport simulation, the simulation results, i.e., cell dose and cell DNA DSB, are used to quantify the cell state transitions. The cell state transition results are used to determine whether cell is killed by radiation or not. If the cell is killed, cell type changes to necrotic. Then the tumor system keeps its evolution path till next irradiation. In this simulation, the total simulation time is 14000 MCS, and after simulation, the tumor growth curve can be obtained.

As shown in Figure 12a, six dose schemes are simulated, and the tumor growth curves of those dose schemes are plotted together for comparison. The tumor growth curves show that the tumor growth rate is reduced with the irradiation. The higher the dose, the more the reduction of growth rate. However, interestingly, we can observe that the tumor growth rate is even higher than the growth rate of no irradiation when the total dose is 5 Gy in 5 fractions. The possible reason for this is that the low dose irradiation kills some cells, which eventually leads to more free space for tumor growth since the dead cells will gradually be eliminated in tumor cluster, and this offsets the space constraint due to contact inhibition for cell proliferation.

**Figure 12:**
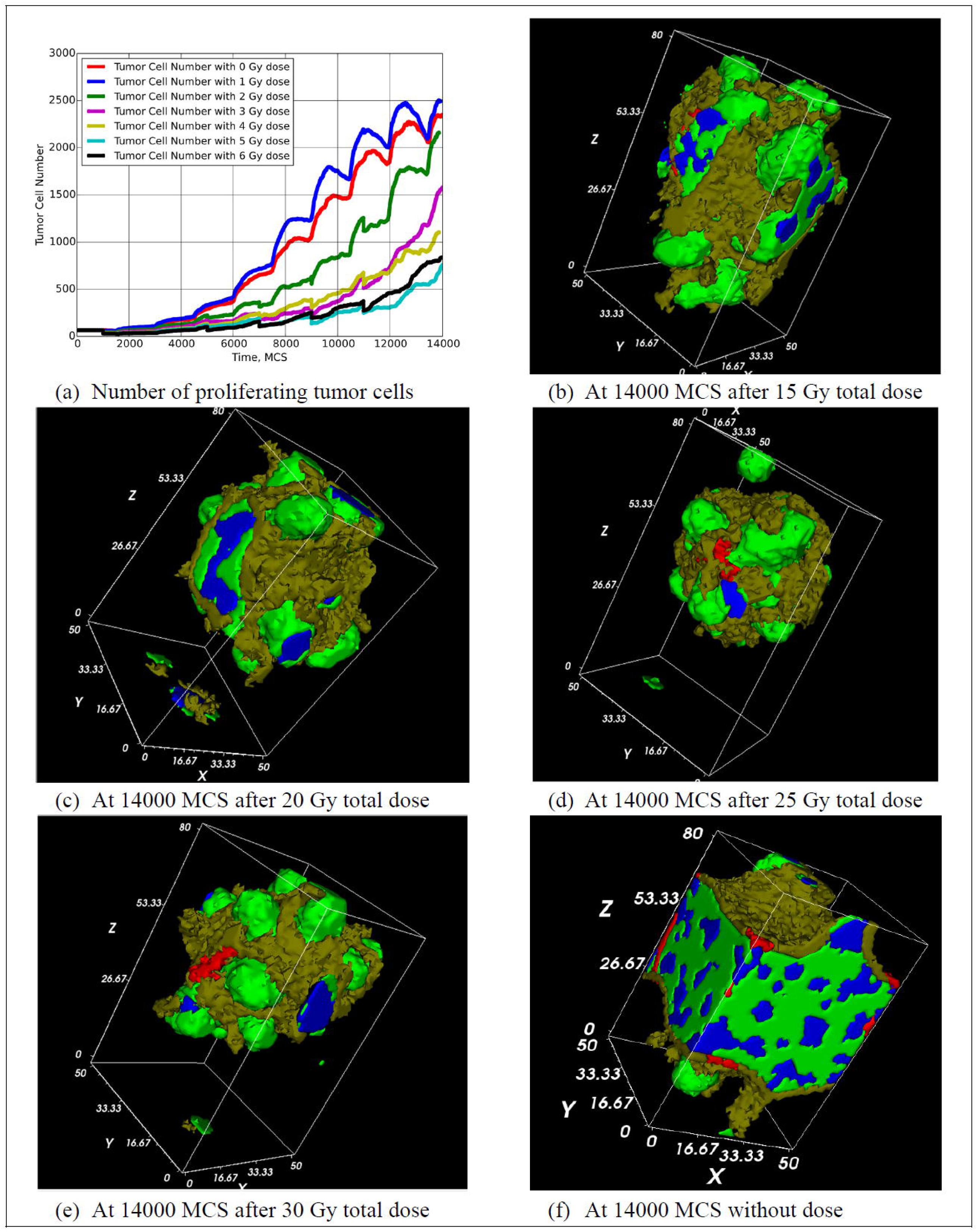
Two-dimensional snapshots of the vascular tumor simulation. Red: vascular cells, grey: neovascular cells, blue: necrotic cells. The spatial unit used in CC3D is pixel, and one pixel equals 4 µm in this work. The dose is delivered in 5 fractions, and the delivering time at: 1000 MCS, 5000 MCS, 7000 MCS, 9000 MCS, and 11000 MCS, respectively. (a) shows the tumor growth curves under different dose levels with respect to tumor evolution time which has MCS as its unit. (b) shows tumor system after irradiation with 15 Gy dose at 14000 MCS rendering in CC3D visualization system. (c), (d), (e), and (f) show the tumor system after irradiation with 20 Gy, 25 Gy, 30 Gy, and 0 Gy, respectively. (f) serves the control compared to the cases with irradiation.

We also can observe that the number proliferating tumor cells drop after each irradiation, but tumors will quickly recover from the cell loss when the dose is relatively low in each fraction, which indicates that tumor cell repopulation can offset the tumor cell loss due to irradiation. Besides analyzing the tumor growth curve, we also can observe the morphological change of tumor after irradiation. Through the CC3D simulation, we can visualize the tumor shape with time. For instance, we predict how the tumor cluster will look like after a series of irradiation. Here, we list the snapshots of tumor cluster under four different dose schemes, as shown in Figure 12.

#### 3.5.3 Tumor Response with Hyperfractionated Dose

It is found that fractionation of the radiation dose produces, in most cases, better tumor control for a given level of normal tissue toxicity than a single large dose. However, treatment with any cytotoxic agent, including radiation, can trigger surviving cells in a tumor to divide faster than before. The critical point is that during the time that tumor is overtly shrinking and regressing, the surviving clonogens are dividing and increasing in number more rapidly than before treatment [44]. There are two separate strategies to cope with this issue, and they are hyperfractionation and accelerated treatment.

In this simulation, we try to simulate a hyperfractionated dose delivering scheme for vascular tumor. We use a single planar microbeam source to irradiate the vascular tumor cluster. Particularly, the tumor cluster size at the first irradiation is chosen to be relatively large, which is used to model a fully-grown tumor. We obtain this “grown” tumor by letting the initial tumor grow without irradiation for 12000 MCS. Two fractionated dose schemes are simulated, the first one is 40 Gy in five fractions, and the second one is 40 Gy in 2 fractions.

The dose delivering time for the first scheme is at: 12000 MCS, 13000 MCS, 14000 MCS, 15000 MCS, and 16000 MCS. The dose delivering time for the second scheme is at 12000 MCS and 16000 MCS. For the first scheme, five doses are delivered within 4000 MCS. As we discussed above, 1 MCS approximately equals 1 minute in this study, so we know that roughly there are two doses per day. So, the first delivering scheme is considered to model a hyperfractionated scheme, and the second scheme is considered to model a hypofractionated scheme.

From the tumor growth curve shown in Figure 13a, we can know that the hyperfractionated scheme performs better in offsetting the tumor cell repopulation after each fractionated dose. On the contrary, the hypofractionated scheme could not offset the tumor cell repopulation. As shown in Figure 13a, after the first fractionated dose in the hypofractionated scheme, the number of tumor cells gradually recover to almost the initial state by the time of second fractionated dose. This result indicates that the hyperfractionated scheme is better for controlling the vascular tumor recovery under the single planar microbeam irradiation.

**Figure 13:**
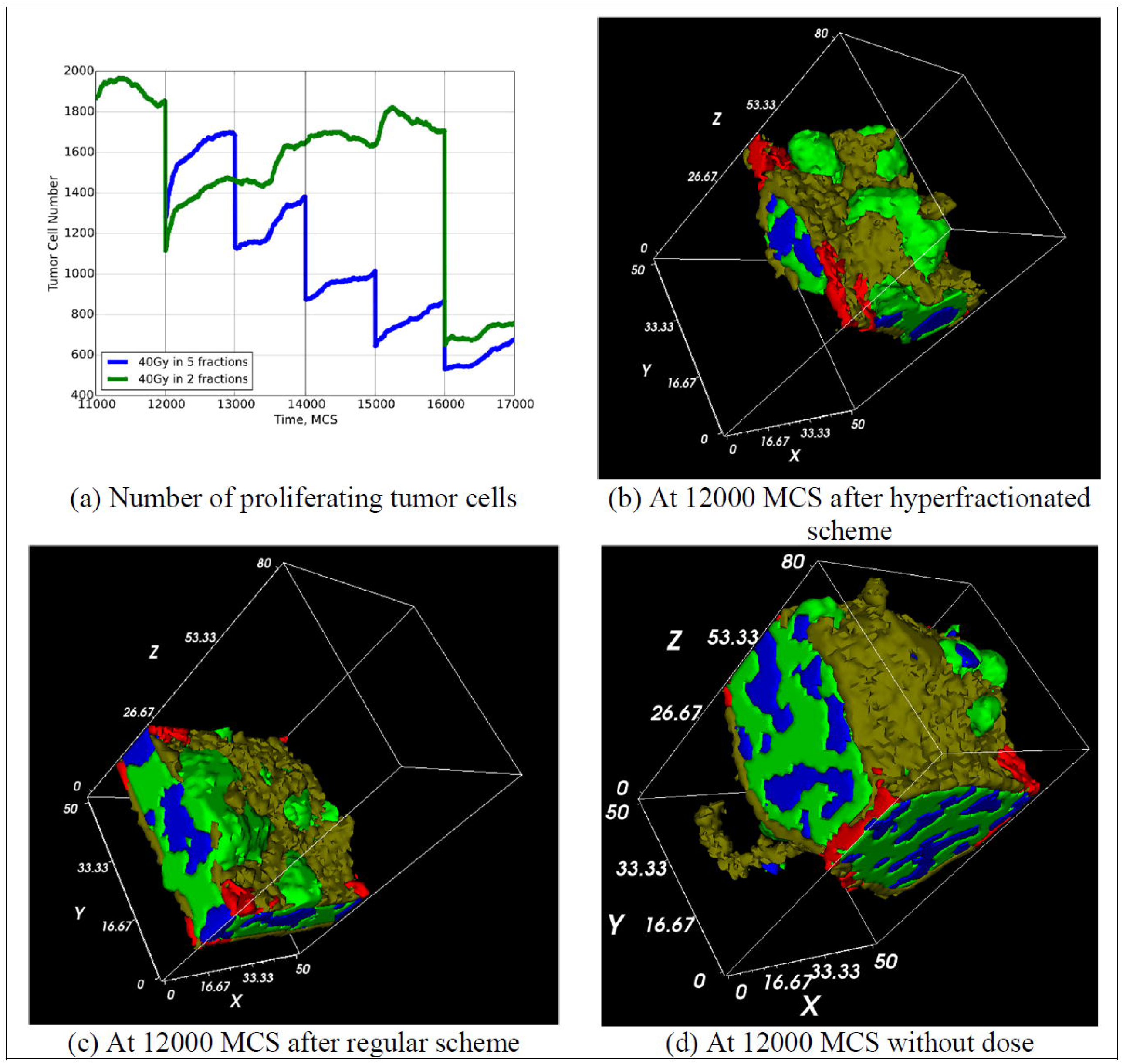
Two-dimensional snapshots of the vascular tumor simulation. Red: vascular cells, grey: neovascular cells, blue: necrotic cells. The spatial unit used in CC3D is pixel, and one pixel equals 4 µm in this work. The hyperfractionated doses are delivered at 12000 MCS, 13000 MCS, 14000 MCS, 15000 MCS, and 16000 MCS. The regular doses are delivered at 12000 MCS and 16000 MCS. (a) shows the tumor growth curve under two different dose deliver schemes. (b) shows the tumor system at 12000 MCS after the hyperfractionated dose deliver scheme. (c) shows the tumor system at 12000 MCS after the regular dose deliver scheme. (d) shows the tumor system at 12000 MCS without irradiation, which serves as control for the cases with irradiation.

The morphological change of tumor after irradiation is shown in Figure 13b and Figure 13c. By comparing the shape of the tumor under those two different dose schemes, we can know that both dose schemes can reduce the tumor volume, and the hyperfractionated scheme reduces more tumor volume.

#### 5.3.4 Tumor Response with Fractionated Dose by MRT

Here, we simulate how the vascular tumor response to the multi-array planar microbeam irradiation. It is worth noting that the MRT in the real clinical application is using a multi-array microbeam source, so this simulation captures the features of a real MRT treatment condition.

In our simulation, there are five planar microbeam sources separated by 200 *μ*m center by center, as shown in Figure 14, and those five sources serve as the multi-array microbeam source. The hyperfractionated dose schemes, i.e., 40 Gy in 5 fractions and 50 Gy in 5 fractions, are selected to deliver the dose to a “grown” vascular tumor. In both schemes, the first irradiation starts at 12000 MCS, and dose delivering time is at: 12000 MCS, 13000 MCS, 14000 MCS, 15000 MCS, and 16000 MCS.

**Figure 14:**
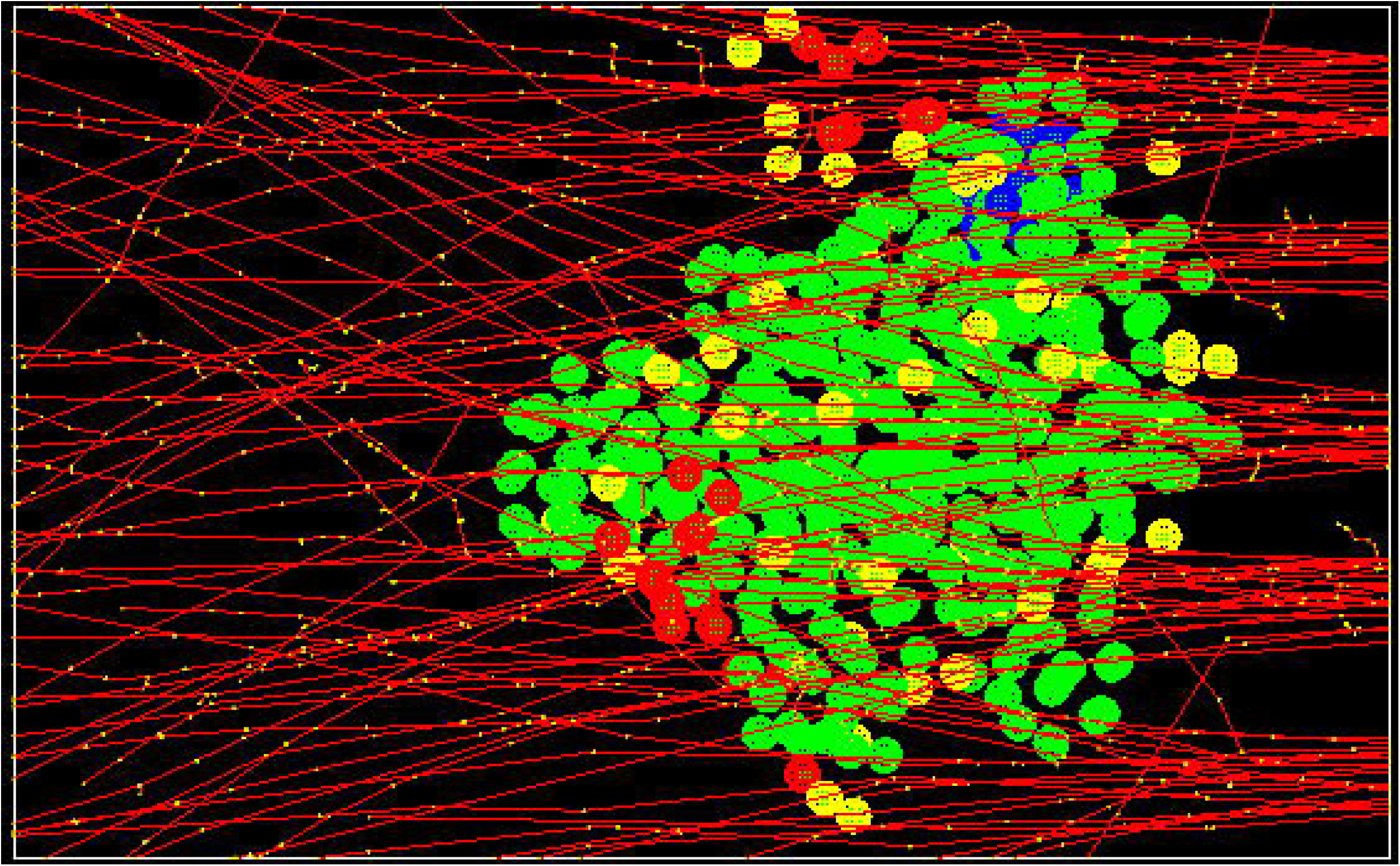
Multi-array planar MRT irradiation on vascular tumor. Five panels of planar microbeams are used to irradiate tumor. In Geant4 platform, the red lines are the tracks generated by electrons, and the yellow points are the energy deposition points after radiation interactions.

From the tumor growth curves shown in Figure 15a, we can see that the hyperfractionated scheme by the multi-array microbeam can substantially reduce the growth of tumor cells. After the whole scheme delivery, the proliferating tumor cells are almost eliminated. We can also observe that the two hyperfractionated scheme performs nearly same in terms of the tumor control since, after the whole scheme, all the tumor cells are almost killed. We can calculate the proliferating tumor cell loss after one fraction of the dose by comparing the tumor cell number before and after the dose delivering. As shown in Figure 15a, there is a steep drop in cell number after the delivery of each dose. By comparing the tumor cell loss after each fraction in both fractionated schemes, we can see that there is no substantial difference in those two schemes, which indicates that there is a saturation of tumor control in terms of the total dose. The morphological change of tumor after multi-array planar microbeam irradiation is shown in Figure 15. We can see that after the multiarray microbeam irradiation, the vascular tumor gradually shrinks its volume.

**Figure 15:**
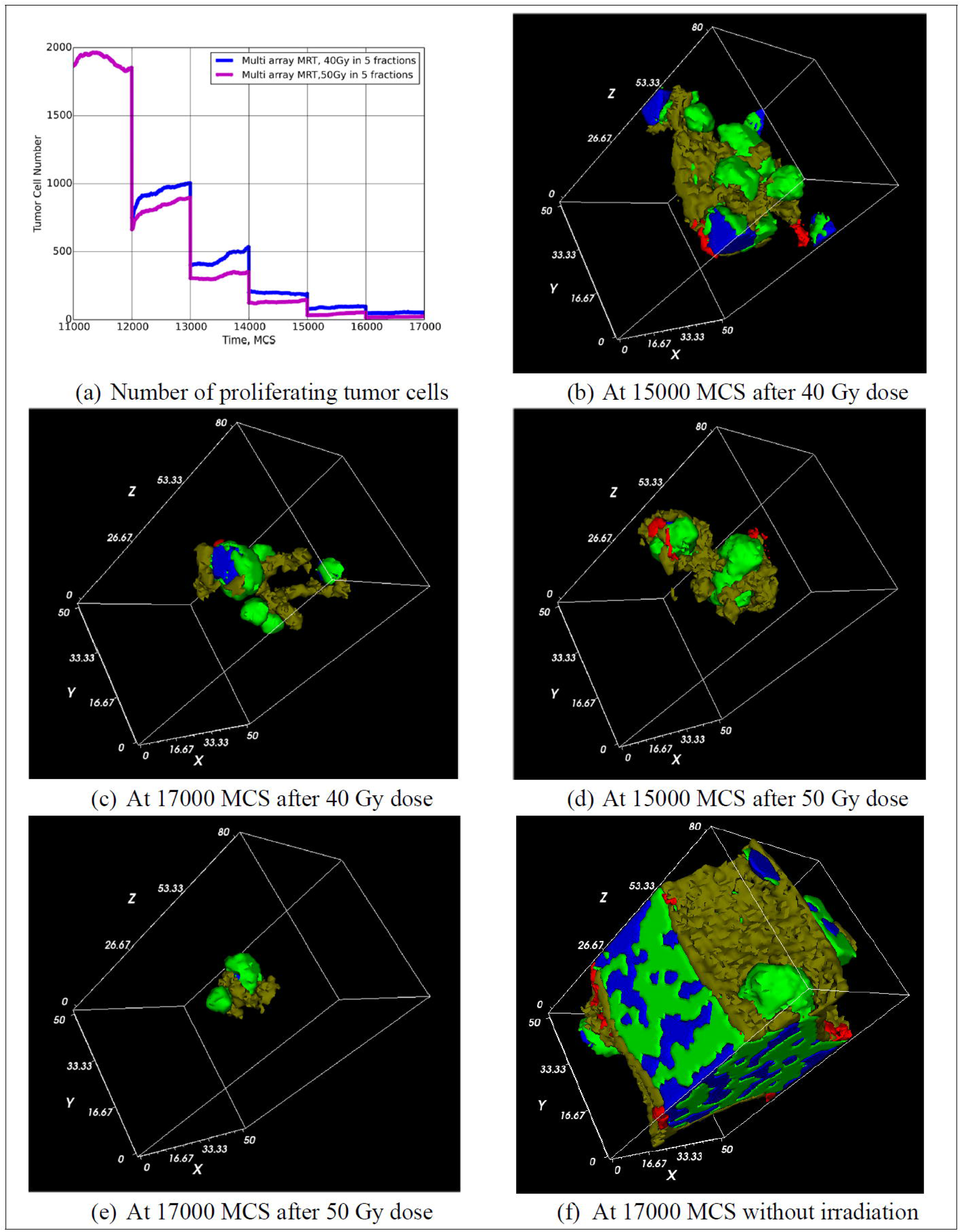
Two-dimensional snapshots of the vascular tumor simulation. Red: vascular cells, grey: neovascular cells, blue: necrotic cells. The spatial unit used in CC3D is pixel, and one pixel equals 4 µm in this work. The doses are delivered in 5 fractions, and the delivering time at: 12000 MCS, 13000 MCS, 14000 MCS, 15000 MCS, and 16000 MCS, respectively. (a) shows the tumor growth curve under two different dose deliver schemes. (b) shows the tumor system at 15000 MCS after irradiation with 40 Gy dose. (c) shows the tumor system at 17000 MCS after irradiation with 40 Gy dose. (d) shows the tumor system at 15000 MCS after irradiation with 50 Gy dose. (e) shows the tumor system at 17000 MCS after irradiation with 50 Gy dose. (f) shows the tumor system at 17000 MCS without irradiation, which serves the control compared to the cases with irradiation.

### 3.6 Future Work

In this work, we simulate the change of cell-level parameters such as target volume after irradiation by integrating Geant4 and CC3D. We linked the macroscopic cell behavior according to the cell state transition determined by DSBs induced by radiation and the glucose concentration, but we did not consider linking the macroscopic cell behaviors to the intracellular response. The CC3D framework allows us to add and solve subcellular reaction-kinetic pathway models inside each cell to simulate the cell-level behavior by using SBML [45]. In future study, we can model the cellular response for molecular concentrations that can steer the behaviors of biological cells by modulating their biochemical machinery. For example, we can incorporate the repair pathways, such as Non-Homologous End Joining (NHEJ) and Homologous Recombination (HR), by applying the well-known models [46][47] about DNA damage repair using the intracellular simulation functionality of CC3D. Oxygen is an important factor moderating cellular radiation response. In this work, we did not explicitly model the oxygen diffusion and reaction process between cells and inside cells. Instead, we expect that the tumor cells are usually hypoxic, and they can secrete a long-diffusion isoform of VEGF-A to induce angiogenesis in hypoxic tissue. By using the diffusion solver of CC3D we can simulate the oxygen diffusion and reaction between cells and inside cells. Then we can include the models [48][49] of oxygen effect for simulating the radiation effect on vascular tumor.

## 4 Conclusions

To the best of our knowledge, this multi-platform simulation is the first piece of research to combine Geant4 and CC3D to implement the coupled cell biology and radiation transport simulation for quantifying cell response after irradiation. In this paper, we focused on introducing this novel framework with relevant technical details and illustrated with a case study. A vascular tumor simulation model based on the developed toolkit was studied in this work. Despite the rescaling of the tumor size, the model produces a range of biologically reasonable morphologies that allow study of how MRT treatment affects the growth rate, size, and morphology of vascular tumors, which will help to design the effective MRT treatment plan in the real clinical trials in the future.

The test simulation shows that the developed model could be potentially used to facilitate the investigation of radiation biology study. This method is appealing because it allows quantitative assessment of radiation damage in each cell, allowing us to further build more a informed model of radiation damage in tissue level. The presented method could be validated by performing a quantitative measurement of tumor spheroid subject to precise radiation dose delivery. In addition to controlling radiation dose, we could also vary the temporal pattern of dose delivery. Those simple experiments could help constrain parameter ranges of our model. We hope that this work will shed light on building a comprehensive mathematical modeling toolkit for computational radiation biology. The simulation parameters of the vascular tumor model are adopted from previously published research papers. By interacting with clinicians and experimentalists, we may gather experimental data to parameterize, calibrate, and validate the model for its clinical applications.

## Acknowledgements

RL acknowledges support from Consortium for Risk Evaluation and Stakeholder Participation (http://www.cresp.org). JAG acknowledges support from National Science Foundation grant NSF 1720625 and National Institutes of Health, National Institute of General Medical Sciences grants U01 GM111243 and R01 GM076692, JAG and MS acknowledge support from National Institutes of Health, National Institute of General Medical Sciences grant R01 GM122424

## Appendix 1: CompuCell3D general introduction

### Appendix 1.1 Effective Energy

Effective energy is the basis for the operation of the CPM model because it determines the behavior of the interactions between cells. The energy is described in two ways: boundary energy or constraints. One of the most important effective-energy terms describes cell adhesion. This is defined by the boundary energy which is calculated using *J* (*τ*(*σ*),*τ*(*σ*^’^)), the boundary energy per unit area between the two cells (*σ,σ*^’^) of types (*τ*(*σ*),*τ*(*σ*^’^)). This is calculated by the sum over all neighboring pixels that form the boundary between two cells:

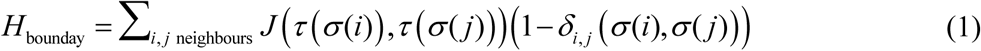

where the delta function *δ*_*i, j*_ (*σ*(*i*),*σ*(*j*)) restricts the boundary energy contribution to cell-cell interfaces. Biologically, higher boundary energies between cells result in greater repulsion between cells, and lower boundary energies result in greater adhesion between cells [1].

The effective energy represents other types of cellular behaviors using constraints. One of the most common use for constraint energies for biological cells in CompuCell3D is to restrict the size of cells to given target values. This is done through two constraints: the volume and the surface area. The volume constraint energy for each cell is calculated by:

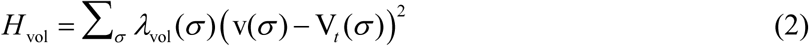

where *λ*_vol_(*σ*) is the inverse compressibility of the cell, *v*(*σ*) is the number of pixels in the cell and *V*_*t*_ (*σ*) is the cell’s target volume in pixels. The constraint is able to define the pressure within the cell as *P* = −2*λ*_vol_ (*v*(*σ*) −*V*_*t*_ (*σ*)). A cell with *v* < *V*_*t*_ has a positive internal pressure, while a cell with *v* > *V*_*t*_ has negative internal pressure.

Since many cells have nearly fixed amounts of cell membrane, the surface area constraint has been introduced as the form:

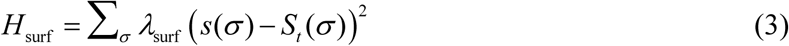

Where *s*(*σ*) is the surface area of cell *σ, S*_*t*_ is its target surface area, and *λ*_surf_ is its inverse membrane compressibility.

Adding the boundary energy and constraint terms together, the basic GGH effective energy is obtained as:

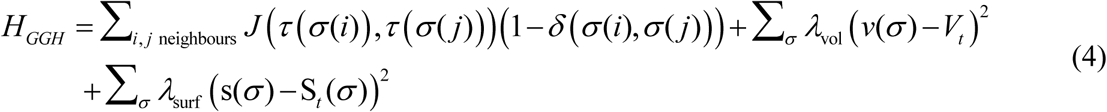

By using equation, the change of effective energy, Δ*H*, could be calculated, which is the basic calculation for determining the dynamics of cell system in CompuCell3D.

### Appendix 1.2 Dynamics

In CompuCell3D, the system evolution is modeled through a dynamic algorithm. A CPM simulation consists of many attempts to copy cell indices between neighboring pixels. In CompuCell3D, the default dynamical algorithm is *modified Metropolis dynamics*. To begin an index copy attempt, a lattice site is randomly chosen to be a target pixel, *i*, and then a neighboring lattice site is chosen to be a source pixel *i*^’^. An attempt will then be made to change the target pixel to the same generalized cell as the source pixel, thereby increasing the volume of the source cell and decreasing the volume of the target cell. If the source and target pixels belong to the same cell (*σ*(*i*) =*σ*(*i*^’^)) then no calculations are necessary.

To determine whether or not an index copy attempt is successful, the following acceptance probability is used:

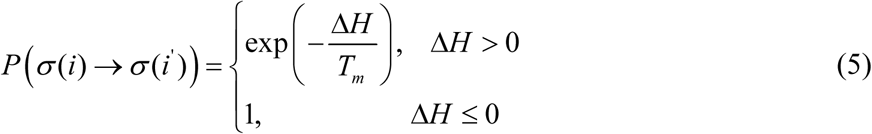

where Δ*H* is the change in effective energy and *T*_*m*_ is a parameter defined to be the effective cell motility, or can be called the amplitude of cell-membrane fluctuations. This Boltzmann acceptance function tells the simulation that if the change in effective energy is less than or equal to zero for an index copy attempt, then the attempt will be successful. However, if the resulting change in effective energy is greater than zero, the attempt may still be successful with the probability exp 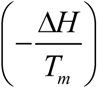.

The Δ*H* could be calculated as

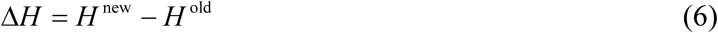

Where *H* ^new^ is the effective energy after pixel copy attempt, and *H* ^old^ is the effective energy before pixel copy attempt. Easily we can know that

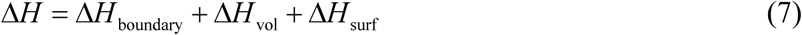

After choosing the source *i* and target *j* pixels (the cell index of the source will overwrite the target pixel if the index copy is accepted), the change of boundary energy is calculated as follows. If the source pixel *i*^’^ belongs to a different generalized cell from the target pixel *i*, when *σ*(*i*) ≠*σ*(*i*^’^), the effective energy *H* decrease by *J* (*τ* (*σ*(*i*)),*τ* (*σ*(*i*^’^))). On the other hand, there is a chance that the source pixel *i*^’^ belongs to a cell, denoting as *σ* (*i*^’’^), which is different from the neighboring cell of the target cell, then *H* increase by *J* (*τ* (*σ*(*i*^’^)),*τ* (*σ*(*i*^’’^))).

The change in volume constraint energy could be calculated as

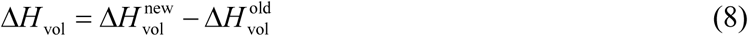

If the pixel copy attempt succeeds, then we can know

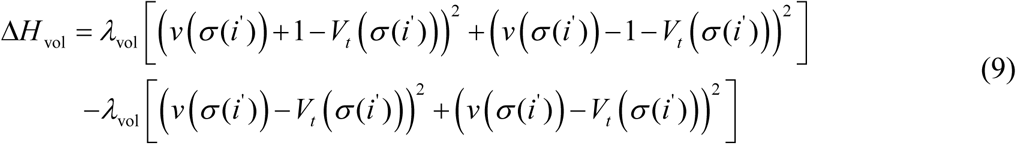

The Δ*H*_surf_ could be calculated similarly as calculating Δ*H*_vol_.

The simulation time in CompuCell3D is measured by Monte Carlo Step (MCS) which is defined as an index-copy attempts, where is the number of sites in the cell lattice. The conversion between MCS and the experimental time depends on the average value of 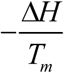 In biologically meaningful simulations, MCS and experimental time are proportional [1].

### Appendix 1.3 Simulation Structure in CompuCell3D - Implementation Details

CompuCell3D uses plugin-based architecture where plugins are understood as software modules that perform a well-defined task within the simulation. For example, modules that calculate effective energy terms are referred to as energy plugins, while modules that monitor events on the cell lattice are called lattice monitors. Thus, every CompuCell3D simulation consists of combining energy plugins, lattice monitors, and additional Python modules that typically perform on-the-fly adjustment of simulation parameters or carry out analyses of the current snapshot of the simulation. Those on-the-fly adjustments are implemented as CompuCell3D steppables, i.e., as a short software module that gets called every Monte Carlo Step. Typically, we implement steppables using Python, but we could use C++ to achieve better computational performance. The Geant4-based extension to CompuCell3D that we developed as a part of this project is coded C++ for optimal speed.

In the section [2.1.2], we explained that CompuCell3D simulation consists of pixel copy attempts where CompuCell3D will evaluate change of effective energy, should the pixel copy be accepted (energy plugins do this task) and if the pixel copy gets accepted CompuCell3D will - using lattice monitors - update attributes of the cells that are affected by the pixel copy (e.g., adjusting volume, surface or center of mass of the affected cells).

The flowchart of the CPM algorithm implemented in CompuCell3D is shown in Figure 1.

**Figure 1:**
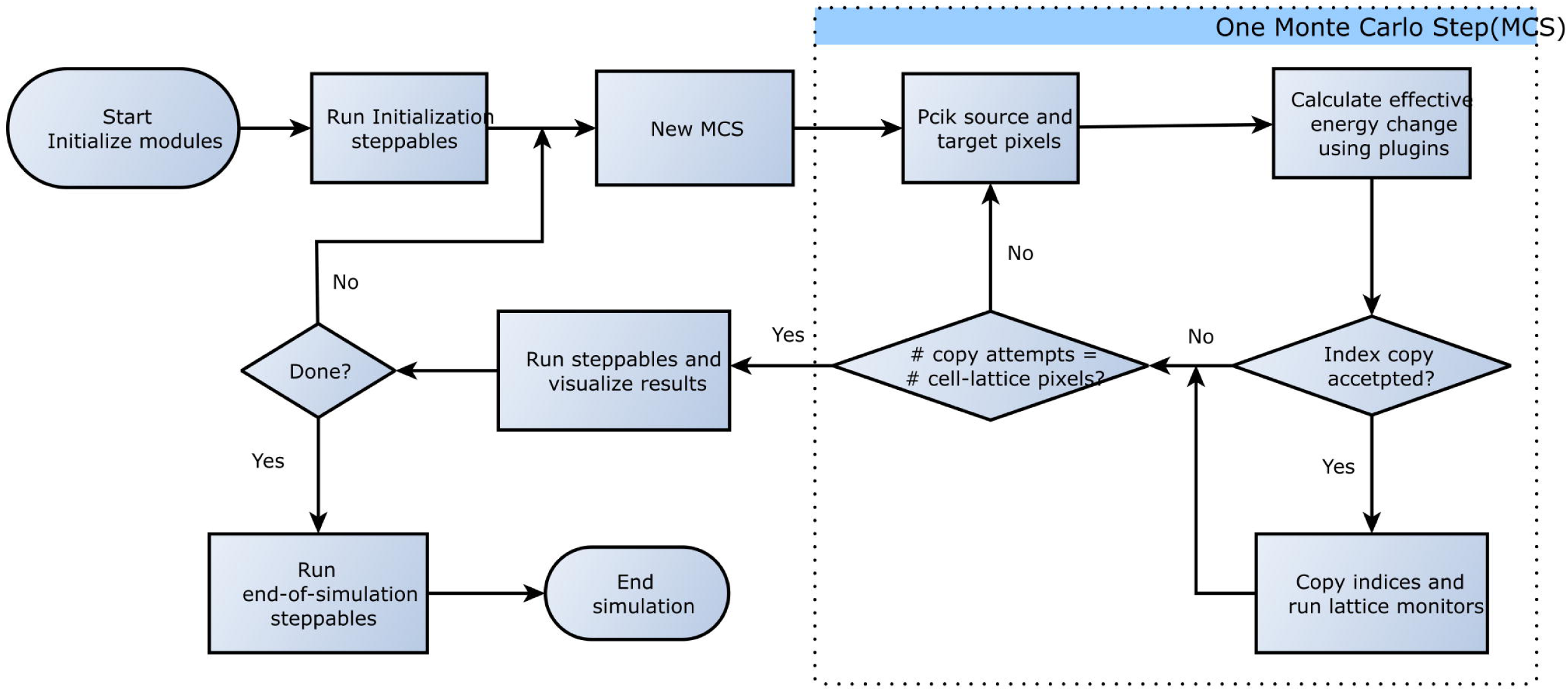
Flowchart of the CPM algorithm as implemented in CompuCell3D [2].

## Appendix 2: RADCELL general introduction

Below we provide a brief description of components of the RADCELL module. The UML diagram of RADCELL is shown in Figure 2. The group of classes in yellow correspond to the standard Geant4 base classes. The core of the radiation transport module is represented by the group of classes in green. The RADCELL provides a natural bridge between CC3D and Geant4, which makes the coupled simulation between Geant4 and CC3D applicable.

### Appendix 2.1 RADCellSimulation

The RADCellSimulation is the class used to control the whole process of radiation transport simulation of cell. It takes care of building the geometry of cell culture, cell visualization, radiation track structure visualization, cell geometry update, etc. The CreateCell() method is used to create a cell line. The SetCellWorld() method is used to build cell culture geometry. These methods are used in the initial phase when we describe cell and tissue geometry in the Geant4 simulation. The UpdateGeometryInitialize() method and UpdateGeometryFinalize() methods are used to update the cell geometry since the cell geometry may change due to cell proliferation or cell death. The EnergyDistributionCalculation() method is used to invoke the Geant4 transport kernel to implement the radiation transport simulation of cells.

### Appendix 2.2 Cell

The Cell class is used to describe one type of cell line. Some attributes are used to describe the characteristics of cell. For instance, cellType indicates the type of cell, cellOrganelle is a vector that stores the cell organelles, and cellShape indicates the shape of cell. The CellConstruct() method is used to construct a cell line. This class is not to be confused with CompuCell3D CellG class. RADCELL’s Cell class is a representation of the CellG class in the RADCELL module, and thus every CellG CompuCell3D object will have matching RADCELL Cell object.

**Figure 2:**
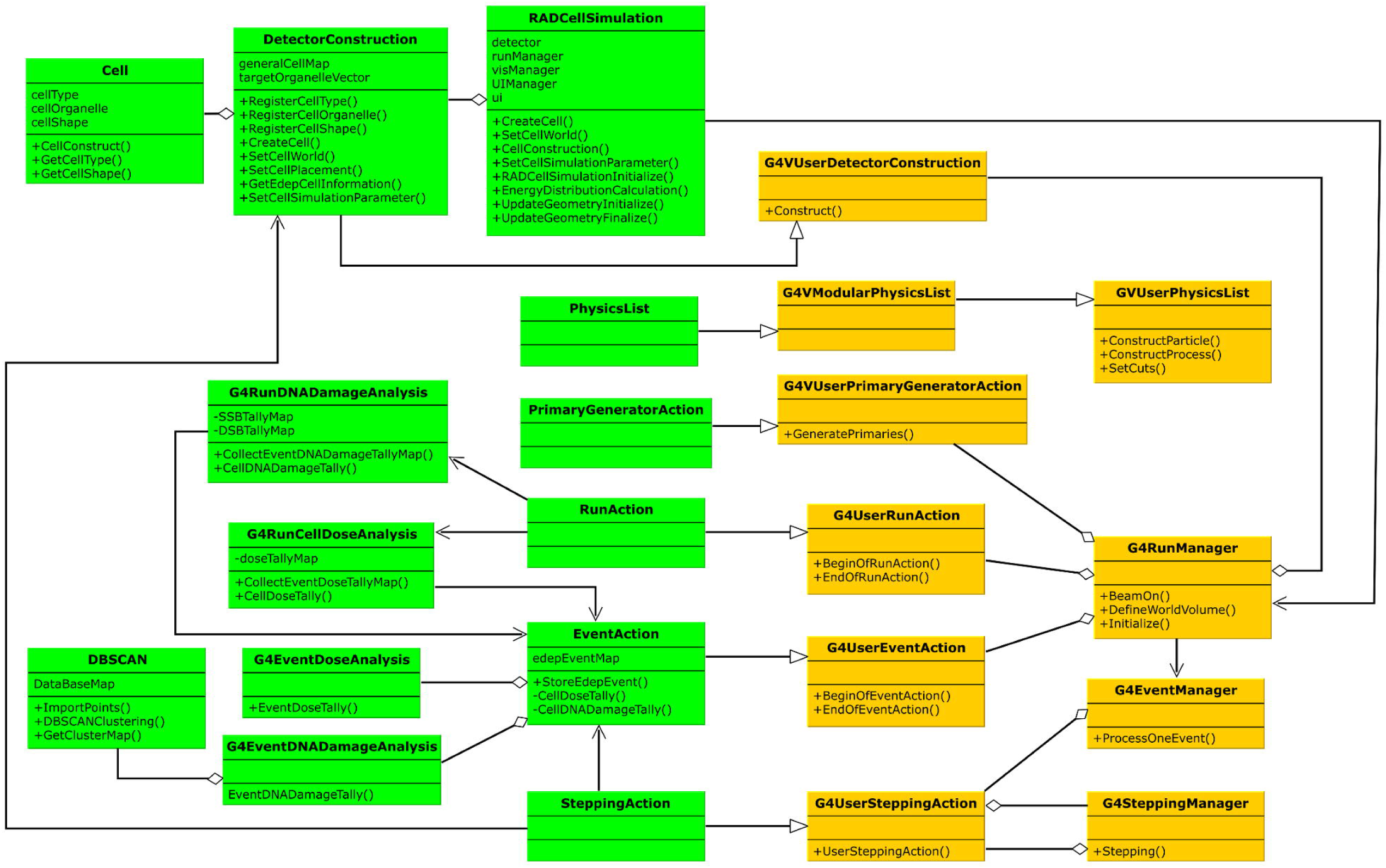
UML diagram of the radiation transport solver: Geant4 base classes (yellow), Geant4 implementation classes (green). (The relationship between bases classes is referred to paper [3]).

### Appendix 2.3 DetectorConstruction

The DetectorConstruction class is used to build simulation geometry. It serves the function of translating the geometry of cell culture to the corresponding geometry in radiation transport simulation. The SetCellWorld() method is used to define the boundaries of simulation geometry. In Geant4, a world is used to define the boundaries of geometry. Usually, we can use a box to define the world where the whole cell culture locates inside. The cells in the cell culture may belong to different cell lines, so before seeding the cell into cell culture, the cell objects of cell lines should be created using the method CreateCell(). The information of cell lines will be stored in a vector generalCellMap. The SetCellPlacement() method is used to seed the cell in cell culture. This method will create an actual physical volume to represent the cells seeded in the cell culture. The SetCellSimulationParameter() method is used to specify the simulation parameters for simulation, such as setting the production cuts of secondary particles and specifying the target logical volume which will apply Geant4-DNA crosssection data. The GetEdepCellInformation() method is used to determine which cell and to which cell organelle the energy deposition point belongs.

### Appendix 2.4 DBSCAN

The DBSCAN is a class used to implement the DBSCAN clustering algorithm [4]. It serves as a clustering kernel for clustering two-dimensional and three-dimensional data. In RADCellSimulation, the energy deposition points will be clustered using the DBSCAN algorithm.

### Appendix 2.5 EventAction

In Geant4, there are two big units of simulation, i.e., Run and Event. An event is the basic unit of simulation in Geant4. The event corresponds to the interactions of one particle from the radiation source with matter (in our case cell). One event contains the whole transport process of one primary particle. The run is the largest unit of simulation. One run consists of a sequence of events. Thus a run corresponds to simulating a time interval in which many events of radiation-matter interactions take place [5]. To control or to keep track of what happens during the event, Geant4 introduces the EventAction class (class directly derived from the base class G4UserEventAction in Geant4 toolkit). The EventAction is the class for defining actions of event. This class has two virtual methods which are invoked by G4EventManager for each event, and they are BeginOfEventAction() and EndOfEventAction(). The EndOfEventAction() method is invoked at the very end of event processing. It is typically used for a simple analysis of the processed event. For scoring simulation results, tally is the process of scoring the parameters of interest; for example, keeping track of energy depositions. In radiation transport simulation, the tally of cell dose and DNA damage for each event will be invoked in this method. In RADCELL, we use CellDoseTally() method to get the cell dose tally of the event and the CellDNADamageTally() to get the cell DNA damage tally of the event.

### Appendix 2.6 RunAction

To keep track of what happens during multiple Geant4 events i.e., during multiple events or radiation-matter interaction events, RADCELL uses RunAction class which is directly derived from the base class G4UserRunAction in Geant4 toolkit. This class has two key methods, BeginOfRunAction() and EndOfRunAction(), defined in the base class G4UserRunAction. The EndOfRunAction() is invoked at the very end of a run, and we use it for storing/printing histogram or manipulating run summary. For example, we implemented final tally of cell dose and DNA damage in this method. For final tallies, Geant4 uses G4RunDoseAnalysis class to do the final tally of cell dose. In G4RunDoseAnalysis, the CollectEventDoseTallyMap()method will collect the dose tally results from all the events, then the CellDoseTally() method will do the final tally of cell dose and statistical error analysis. Similarly, G4RunDNADamageAnalysis class is created to do the final tally of DNA damage and statistical error analysis.

In G4RunDNADamageAnalysis, the CollectEventDNADamageTallyMap() method will collect the DNA damage tally results from all the events, then the DNADamageTally() method will do the final tally of cell DNA damage. When the run is finished, one object of G4RunDoseAnalysis will be instantiated inside the EndOfRunAction() method to get the final dose tally results of a run. Similarly, one object of G4RunDNADamageAnalysis will be instantiated inside the EndOfRunAction() method to get the final DNA damage tally of run. The calculation process of dose tally and DNA damage tally is shown in Figure 3.

**Figure 3:**
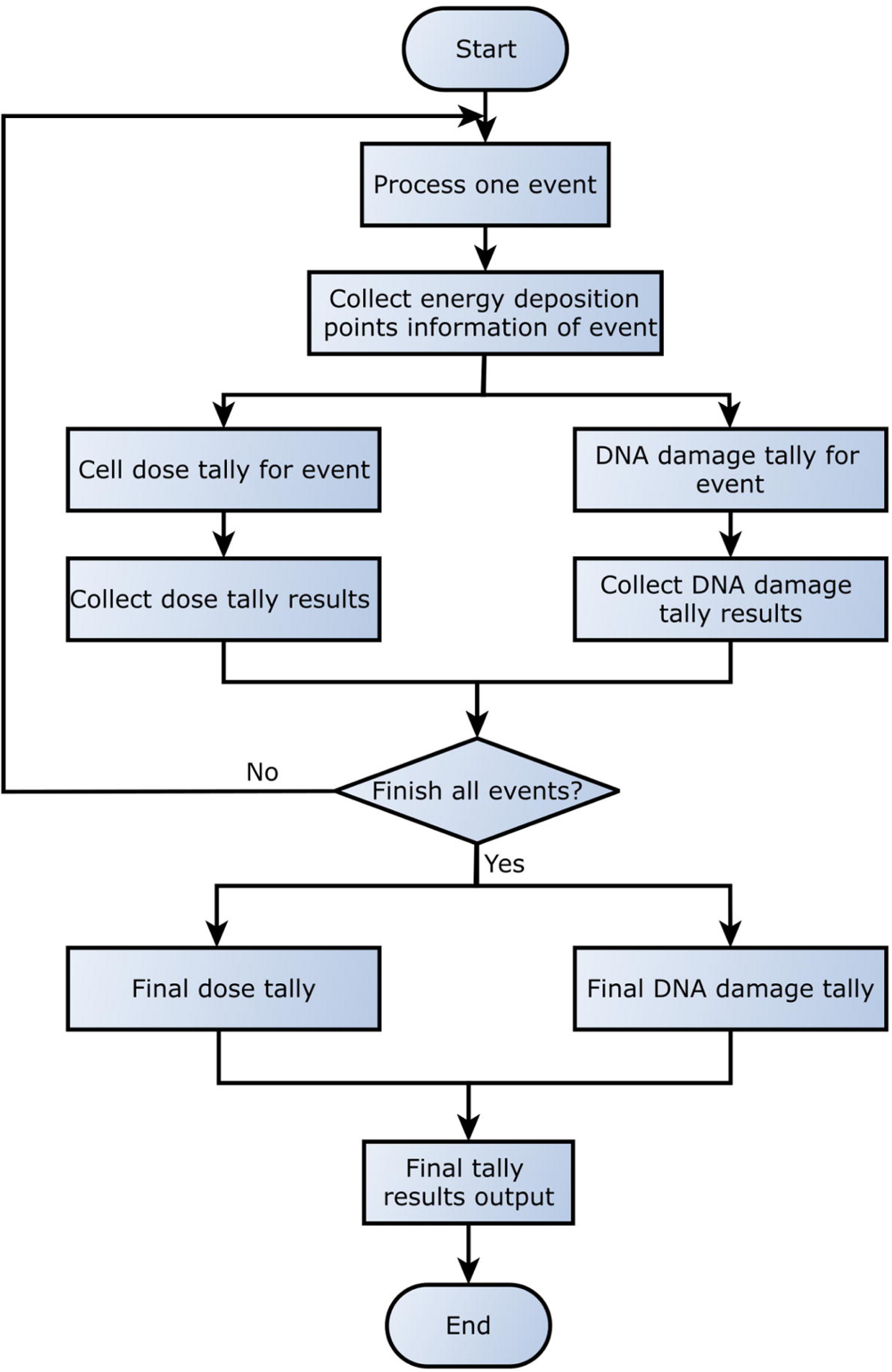
The process of implementation cell dose and DNA damage tally of cells.

### Appendix 2.7 PhysicsList

The way Geant4 simulates interaction of radiation with matter is by keeping track of various physical mechanisms in which the radiation will occur. Specifying which interactions are relevant for our simulation takes place in the PhysicsList class which is derived, in large part, from the microdosimetry example included in the Geant4 toolkit. This class activates a full suite of Geant4-DNA specific physics included in Geant4. Physics processes and models in Geant4 are enabled by region which is known in program as G4Region. In G4Region, Geant4-DNA physics are enabled.

### Appendix 2.8 PrimaryGeneratorAction

Once we defined cells in the Geant4 simulation along with all the infrastructure necessary to keep track of radiation-induced effects, we need to specify radiation source. We use so called general physical source (GPS) within radiation transport solver to describe properties of the radiation source. The GPS allows users to define the source according to the efficient built-in accommodations for source location distributions, energy distributions, angular distributions, and other features.

### Appendix 2.9 SteppingAction

PhysicsList defines many types of interactions that may happen when a single event of radiation-matter interaction happens. Geant4 will simulate those types of interaction sequentially, and it uses SteppingAction class to allow users to keep track of what happens when a single type of interaction takes place. The SteppingAction is derived, in large part, from the microdosimetry example included in the Geant4 toolkit. SteppingAction inherits the UserSteppingAction() method from the base class G4UserSteppingAction. When the UserSteppingAction() is invoked, we keep track of the information of energy deposition and store it in the EventAction object.

## Appendix 3: Cell State Transition Model

Mathematically, we denote the possible cell states as a set {*S*_1_, *S*_2_, *S*_3_} [6]. *S*_1_ is corresponding to healthy state, *S*_2_ is corresponding to arrested state, and *S*_3_ is corresponding to dead state. To model cell state transitions after radiation exposure, we define a *state energy* to quantify the radiation-induced damage to each cell. Each cell state *S* has a state energy *E*. In this work, the cell state energy is calculated by

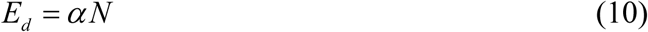

where *α* is a constant and *N* is the number of DSBs produced by direction radiation hitting the cell. The DSB is the most lethal damage type to cell, and here we aggregate all direct damage into an effective DSB number.

When a cell experiences irradiation directly or absorbs bystander signal, the cell may have DNA damage and its state energy increases. When DNA damage is repaired, the state energy decreases. When state energy increases, cell has more potential to have cell state transition from healthy state to arrested or dead state. On the other hand, when state energy decreases, cell has more potential to have cell state transition to healthy state. We introduce another term, *external perturbation energy*, Δ*E*, to quantify the net change of the state energy under the external influence. Suppose the initial state energy is *E*_*i*_, after cell absorbing external perturbation energy, the state energy will change to

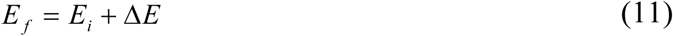

Where *E*_*f*_ is the final state energy.

We introduce a cell state energy distribution for each cell state, and we propose that each state energy follows a normal distribution. The state energy distribution of state 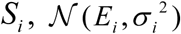, is written as:

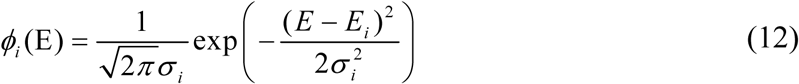

where *E*_*i*_ is the mean state energy of state *S*_*i*_, and *σ* is the standard deviation of state energy of state *S*_*i*_. The *E*_*i*_ is taken as the most feasible state energy for state *S*_*i*_, and *σ* is a measure of the fluctuation of state energy distribution.

If cell gets direct DSB damage, then cell goes through instantaneous transition. The instantaneous transition takes place in the current time step when cell gets direct DSB. The cell state transition probability from state *S*_*i*_ to *S* _*j*_ is calculated as:

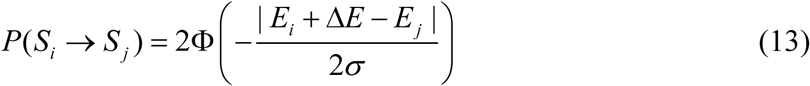

where *σ* is the standard deviation assuming that state energy follows a Gaussian distribution.

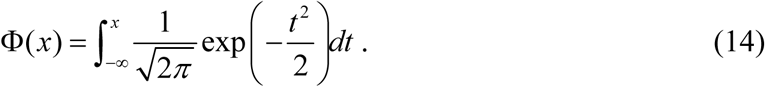

A detailed description of the development of the algorithm for quantifying the of cell state transition after cell irradiation could be seen in our previous work [6]

## Appendix 4: Cell Models Used in CC3D Modeling

### Appendix 4.1 Cell Interactions

Following basic assumptions of the CPM model [7], we simulate cell interactions by specifying effective energies associated with particular cell behaviors. To represent cell-cell adhesion, we use a boundary energy term defined as follows:

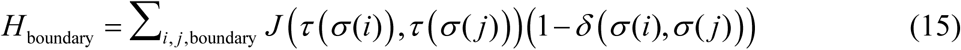

Where *J* (*τ* (*σ* (*i*)),*τ* (*σ* (*j*))) is the boundary energy per unit area between the two cells (*σ,σ*^’^) of types (*τ* (*σ*),*τ* (*σ* ^’^)), the sum is over all neighboring pairs of cell lattice sites *i* and *j*, and *δ* is the Kronecker delta function.

For the purpose of simple nomenclature, we denote boundary energy *J* (*τ* (*σ* (*i*)),*τ* (*σ* (*j*))) between two different cell as *J*_*i, j*_, and we write all the boundary energy pairs into a matrix *J*. In general, *J* matrix reflects how strong the adhesive interactions between cells are. The general rule for determining *J* could be referred to [7][8]. In this study, the default boundary energy matrix, *J* is taken directly from [9]. And it is as

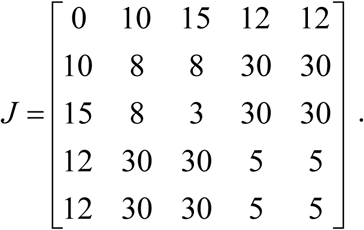

Indices *i* ∈{0,1, 2, 3, 4} correspond to ECM cell, P cell, N cell, EC cell, and NV cell respectively. Thus, for example, *J*_0,0_ corresponds to the boundary energy between ECM and ECM cells, and *J*_2,3_ corresponds to the boundary energy between N cell and EC cell.

In addition to adhesive interactions, we define two additional effective energy terms that implement *cell-volume* constraint:

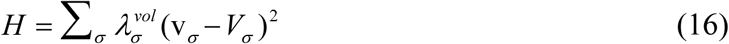

where *v*_*σ*_ denote a generalized cell’s instantaneous volume or instantaneous surface area, and 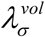 denote the inverse compressibility of the cell.

By combining the boundary energy and constraint energies, we can obtain the basic CPM effective energy as:

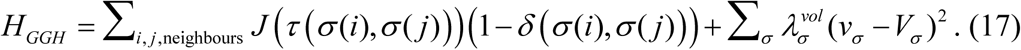

The dynamics of the model is captured by changes in the Hamiltonian, Δ*H*, during each voxel copy attempts according to the rules of the CPM model [7], and it is as:

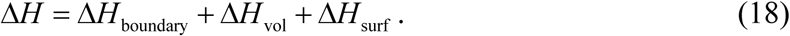

### Appendix 4.2 Cell Growth and Mitosis

Tumor growth *in vivo* depends on the levels of multiple diffusing substances. In this study, we assume that glucose is the primary growth-limiting substance for tumor cell and include a diffusing field representing glucose. The L-VEGF is the primary growth-limiting substance for neovascular cell and includes a diffusion field representing L-VEGF. The concentration rate of glucose could be described by the following reaction-diffusion:

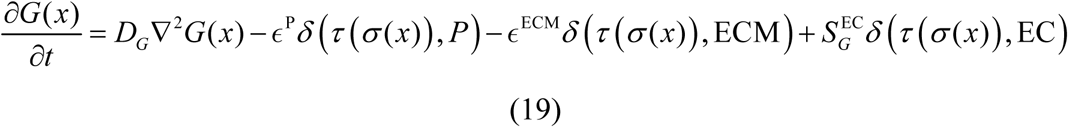

Where *D*_*G*_ is the diffusion constant of glucose, *ϵ*^P^ and *ϵ*^ECM^ are the glucose uptake rate of tumor and ECM respectively, *S* ^EC^ is the secretion rate of glucose of vascular cell. The *δ* (*τ* (*σ* (*x*)), P) means cell *σ* (*x*) is tumor cell, *δ* (*τ* (*σ* (*x*)), ECM) means cell *σ* (*x*) is ECM, and *δ* (*τ* (*σ* (*x*)), EC) means cell *σ* (*x*) is vascular cell.

Similarly, the reaction-diffusion equation that governs VEGF field evolution can be written as:

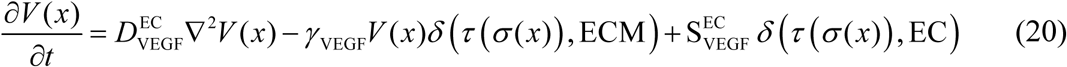

where *δ* (*τ* (*σ* (*x*)), EC) = 1 for inside endothelial cells and 0 elsewhere, and *δ* (*τ* (*σ* (*x*)), EC) = 1 for inside medium and 0 elsewhere. The 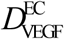 is the diffusion constant, *γ*_VEGF_ is the decay constant, 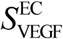 is the secretion rate. The above reaction-diffusion equations governing the glucose and L-VEGF are solved using CC3D’s built-in PDE Solvers [10].

To model tumor growth, we assume that target volume of a tumor cell evolves according to the following relation (that mathematically looks identical to Michaelis-Menten expression from enzyme kinetic):

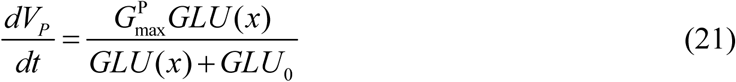

where *GLU* (x) is the GLU concentration at the cell’s center-of-mass (COM), and *GLU*_0_ is the concentration at which the growth rate is half of its maximum. The 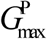 is the maximum growth of tumor cell.

To account for contact-inhibited growth of NV cells, when their common surface area with other EC and NV cells is less than a threshold, we increase their target volume according to:

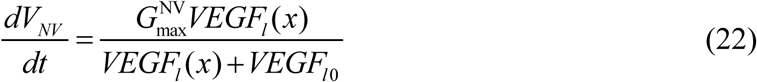

where *VEGF*_*l*_ (*x*) is the concentration of L-VEGF at the cell’s COM, *VEGF*_*l*0_ is the concentration at which the growth rate is half of its maximum, and 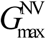 is the maximum growth rate for NV cells. To simulate contact inhibition of vascular cell proliferation we calculate the common surface area between each NV cell and its neighboring NV or EC cells. If the common surface area is smaller than 45, then we increase its target volume. When the volume of NV and P cells reach a doubling volume, we divide them along random axis [9].

## Appendix 5: Cell State Transition

We define the cell state transition rule by glucose concentration as:

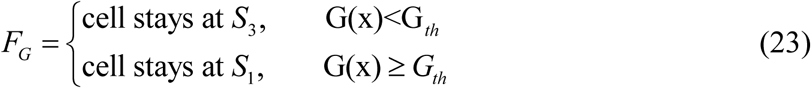

where *G*_*th*_ is the threshold of glucose concentration below which cell dies.

For quantifying the cell state transition after irradiation, we only consider the direct effect of irradiation. The DSB number of cells will be calculated firstly, then the state transition probability from healthy state to dead state is calculated by

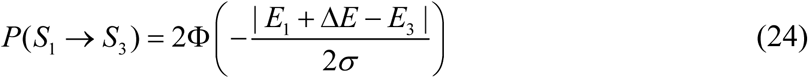

We take *P*(*S*_1_ → *S*_3_) as the cell death probability by radiation effect after irradiation. Then the cell state transition rule by radiation is defined as

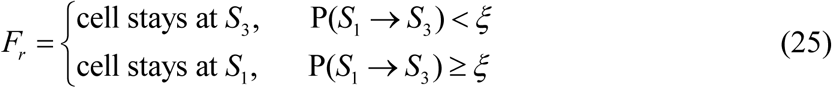

where *ξ* is the random number and *ξ* ∈ [0,1].

Then the cell state transition rule considering the nutrient and radiation conditions is as:

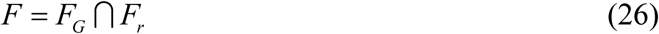

The cell state is updated step-wisely according to the cell state transition rule defined in equation (26).

## Appendix 6: Simulation Parameters

Below we list all simulation parameters (including those from CC3D as well as those from RADCELL model components). These parameters are listed in Table 1.

**Table 1:**
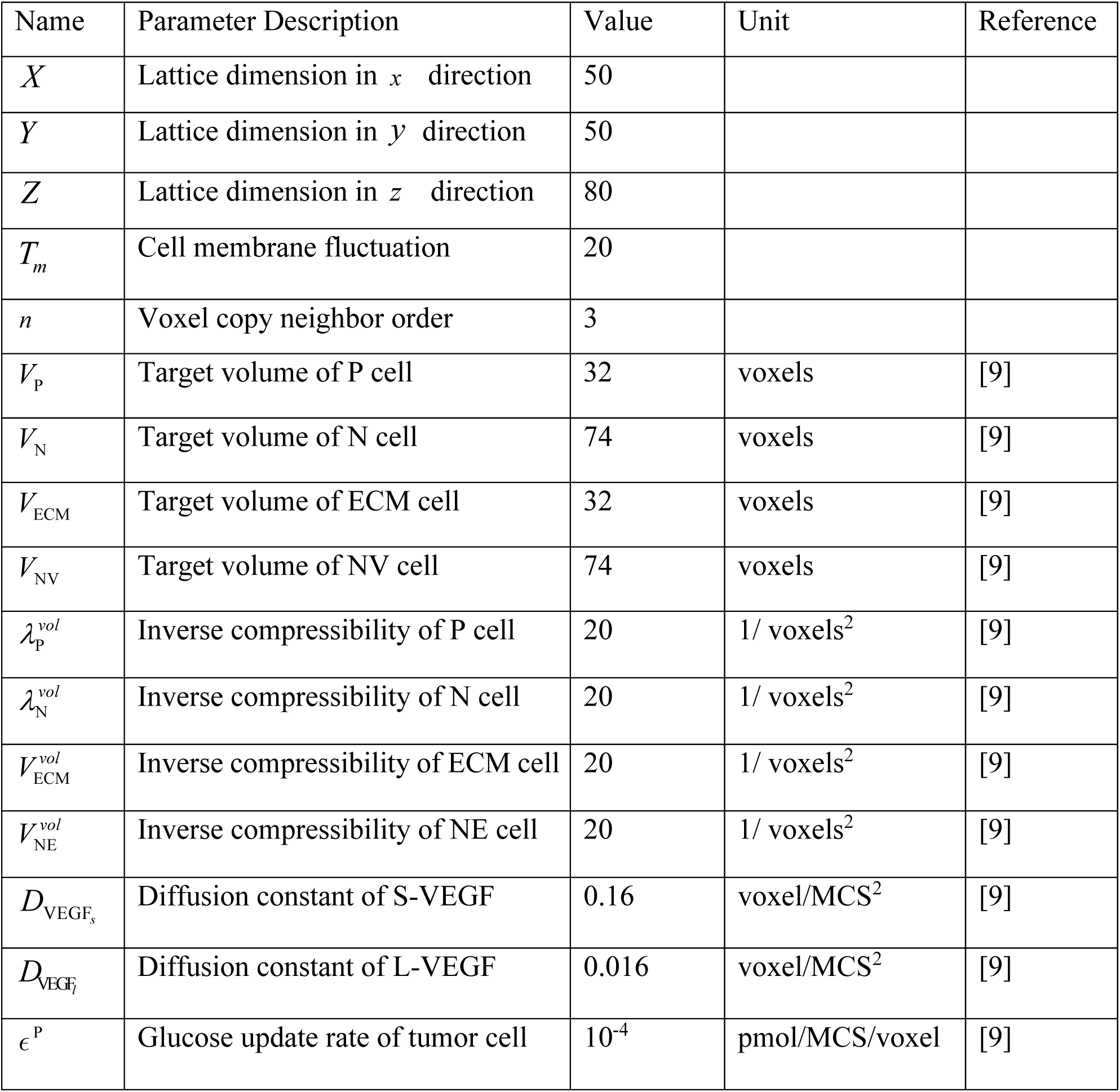
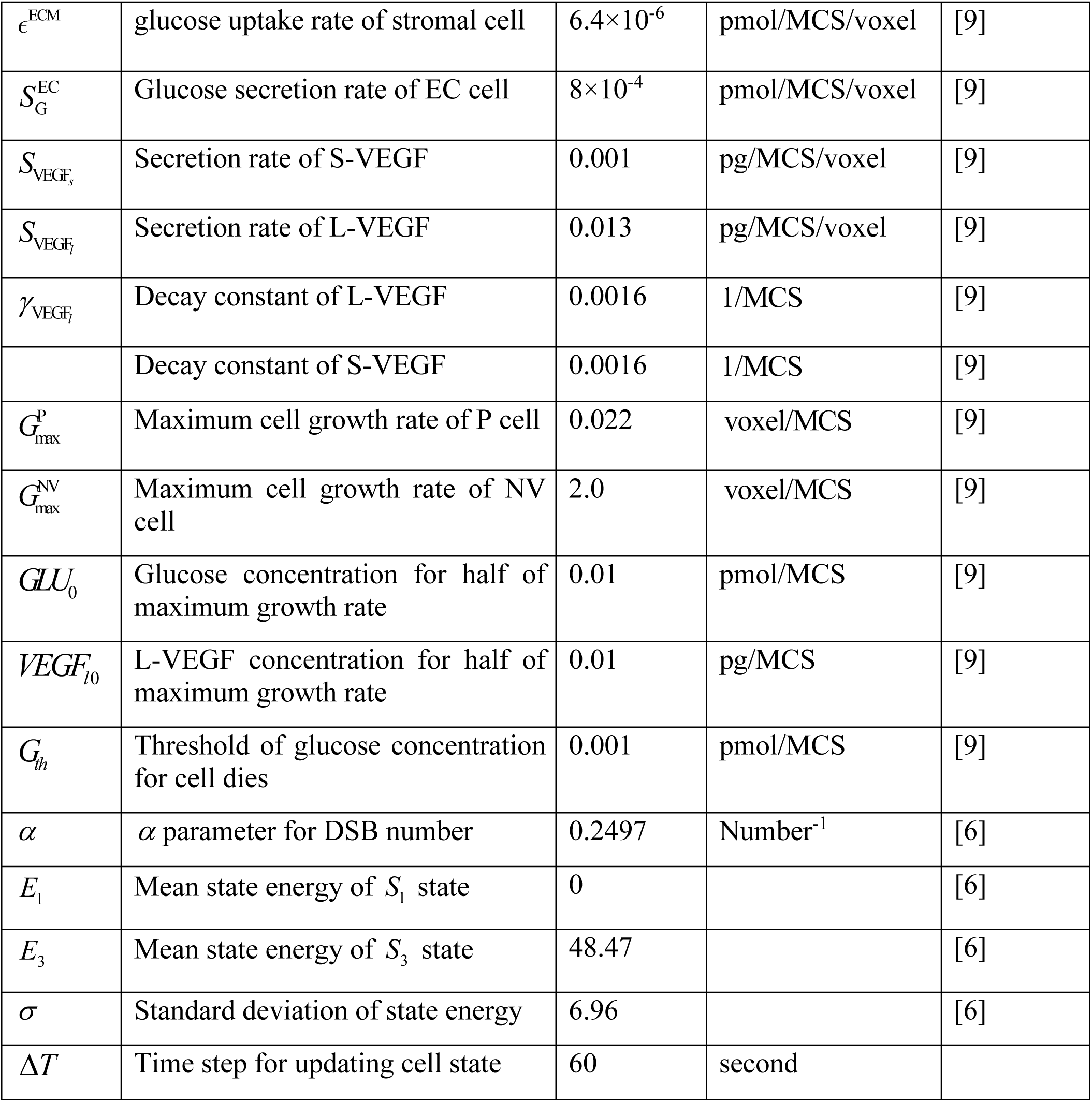
Simulation parameters for coupled simulation

## Notes

### Competing Interest Statement

The authors have declared no competing interest.

